# Modular architecture confers robustness to damage and facilitates recovery in spiking neural networks modeling *in vitro* neurons

**DOI:** 10.1101/2025.02.04.636401

**Authors:** Takuma Sumi, Akke Mats Houben, Hideaki Yamamoto, Hideyuki Kato, Yuichi Katori, Jordi Soriano, Ayumi Hirano-Iwata

## Abstract

Impaired brain function is restored following injury through dynamic processes that involve synaptic plasticity. This restoration is supported by the brain’s inherent modular organization, which promotes functional separation and redundancy. However, it remains unclear how the modular structure interacts with synaptic plasticity, most notably in the form of spike-timing-dependent plasticity (STDP), to define the damage response and recovery efficiency. In this work, we numerically modeled the response and recovery to damage of a neuronal network *in vitro* bearing a modular structure. Consistent with the *in vitro* observations, the *in silico* numerical model effectively captured the decline and subsequent recovery of spontaneous activity following the injury. We revealed that the modular structure confers robustness to injury, minimizes the decrease in neuronal activity, and promotes recovery via STDP. Finally, using the reservoir computing framework, we show that information representation in the neuronal network improves with the recovery of synchronous activity. Our work provides an experimental-numerical platform for predicting recovery in damaged neuronal networks and may help developing effective models for brain injury.

## Introduction

Brain injury impairs crucial brain functions, including cognition, memory, and higher-level executive functions (Walker and Tesco, 2013). Although injuries may temporarily affect large areas of the brain, these functions are often restored spontaneously or through rehabilitation within a few months (Christensen et al., 2008; Chen et al., 2010; Nudo, 2013; Murata et al., 2015; Edlow et al., 2021). Restoration of neuronal activity is central to the recovery of brain function, a process that has been extensively investigated *in vivo* (Fukui et al., 2020) and *in vitro* (Teller et al., 2020; Ayasreh et al., 2022). This restoration is believed to be mediated by network-wide reorganization of functional connections through synaptic plasticity. Clinical and computational studies have indicated that spike-timing-dependent plasticity (STDP), a form of Hebbian synaptic plasticity, is a potential mechanism for restoring damaged neuronal networks (Bunday and Perez, 2012; Gabrieli et al., 2020; O’Neill et al., 2023). Taken together, these findings suggest that synaptic plasticity restores neuronal activity, which in turn leads to the recovery of higher brain function.

In addition to plasticity, the inherent network topological traits are highly involved in injury response and recovery (Arnemann et al., 2015; Boroda et al., 2021). Recent brain connectome analyses have revealed that the brain network evolutionarily conserved a modular structure in which densely connected neuronal populations (modules) are relatively sparsely connected to other modules (Meunier et al., 2010; Lee et al., 2016; Lynn and Bassett, 2019). The coexistence of highly clustered modules and shortcuts between modules facilitates redundancy, ensuring resilience to failure by diversifying information flow, thereby granting robustness to network functions. For example, animal studies have shown that modularity of the frontal cortex increases task feasibility under partial perturbations (Chen et al., 2021). Furthermore, modularity is advantageous for functional recovery, as patients with higher modularity of the cortex exhibit greater improvements in executive function during cognitive training after brain injury (Arnemann et al., 2015). However, it remains unclear how the underlying modular structure of the neuronal network concords with synaptic plasticity, prominently STDP, to lay out efficient damage response and recovery.

In this study, we conducted experiments on cultured neurons and a spiking neural network (SNN) model, both of which were designed with modular organization as the central feature, to examine their response to damage. We created modular neuronal cultures using topographically modulated substrates shaped as parallel tracks, which effectively guided and constrained connectivity along tracks (Montalà-Flaquer et al., 2022) and later inflicted damage using a scalpel. Immediately following injury, the rate of spontaneous neuronal activity was substantially reduced but recovered to its original level after 24 h. Next, an SNN model was devised, and its parameters were manually adjusted to match experimental observations, resulting in a similar reduction and recovery of network activity after damage. The constructed SNN was then used to investigate the effects of damage at different module locations, and to examine the response to damage in networks without a modular structure. Network analysis of the damaged networks suggested that the underlying modular organization reduced the alteration of community structures due to damage, leading to faster recovery than non-modular networks. Finally, the recovery of neuronal activity was conceptually linked to the recovery of brain function using a spoken digits classification task within a reservoir computing framework. Our results highlight the interplay between network architecture and plasticity in facilitating the recovery of activity and (simple) functionality after damage, offering new insights into recovery from brain injury.

## Materials and Methods

### Fabrication of engineered topographical substrates

Polydimethylsiloxane (PDMS) topographical substrates (Montalà-Flaquer et al., 2022) were used to fabricate cultured neuronal networks with modular characteristics. The substrates were prepared using a specially designed mold made of fiberglass and copper (2CI Circuitos Impresos, Spain) to form a two-layer structure. The bottom layer consisted of a uniform fiberglass sheet 2 mm thick, while the top layer contained copper motifs 70 μm high, shaped as parallel stripes 300 μm wide, separated by 300 μm fiberglass gaps, extending the entire length of the mold. In the remainder of this paper, we refer to this pattern as “tracks.”

PDMS (Sylgard 184, Ellsworth Adhesives) was poured onto the fiberglass-copper molds as a mixture of 90% base and 10% curing agent, and cured at 90°C for 2 h. The PDMS was then gently peeled off to form a two-level substrate, which was the negative of the original mold, with PDMS valleys corresponding to copper imprints and crevices to the fiberglass, respectively. The PDMS sheet was perforated using a stainless-steel punch (Bahco 400.003.020), resulting in disks 6 mm in diameter and 1 mm in thickness. The topographical “tracks” pattern on the surface contained 300 μm wide modulations extending across the entire disk. Conceptually, each track gave rise to a module in which neurons are strongly connected along the tracks and weakly connected across them. Before culturing, the PDMS disks were cleaned with ethanol, dried, and mounted on clean coverslips (#1 Marienfeld Superior; 13 mm diameter). One or two PDMS disks were placed on each coverslip, and the PDMS-glass assembly was sterilized in an autoclave (Selecta 4002515). Sterilization increased the bonding between the PDMS and the glass surface such that the PDMS remained attached to the glass throughout the lifespan of the culture.

### Cell culture and GCaMP6s viral transduction

Primary neurons derived from rat embryonic cortices on days 18–19 were used in all experiments. Animal experiments and tissue manipulations were conducted following the approval order B-RP-094/15-7125 from the Ethics Committee for Animal Experimentation of the University of Barcelona (CEEA-UB). Rats were provided by the Animal Farm of the University of Barcelona. Brain dissection was performed in ice-cold L-15 medium (Gibco), and cortical tissue was mechanically dissociated by pipetting. Neurons were then transferred to a ‘plating medium’ [90% Eagle’s minimum essential medium (MEM, Invitrogen), 5% horse serum (HS, Invitrogen), 5% bovine calf serum (Invitrogen), 1 μL/mL B27 (Sigma)]. Before seeding the neurons, the PDMS topographical surfaces were coated overnight with a solution of 20 μg/mL poly-L-lysine (PLL; Sigma-Aldrich) in a borate buffer. Neurons were seeded onto these surfaces to create cultured neuronal networks at a density of approximately 400 neurons/mm^2^. On day *in vitro* (DIV) 1, neurons were transduced with the genetically encoded fluorescence calcium indicator GCaMP6s (AAV9.Syn.GCaMP6s.WPRE.SV40, Addgene) under the Synapsin 1 promoter, so that only mature neurons (and not other cells such as glia) expressed the fluorescence indicator. On DIV 5, the ‘plating medium’ was replaced with a ‘changing medium’ (90% MEM, 10% HS, 0.5% 5-Fluoro-2-deoxyuridine) to limit glial growth. On DIV 8, the medium was switched to ‘final medium’ (90% MEM and 10% HS), which was periodically refreshed every 3 days. Cultured cells were incubated at 37°C, 5% CO_2_, and 95% humidity (Memert INCO2-246).

### Monitoring of neuronal activity

Wide-field calcium imaging of the PDMS topographical cultures was performed using an inverted fluorescence microscope (Zeiss Axiovert 25C, Zeiss GmbH) equipped with a high-speed camera (Hamamatsu Orca Flash 4.0v3, Hamamatsu Photonics) and a fluorescent light source (mercury vapor arc lamp, Osram GmbH) at 12–13 DIV. The fluorescence image series was recorded at 50 frames/s, 8-bit grayscale levels, with an image size of 1,024 × 1,024 pixels. The recordings covered a field of view of 7.1 × 7.1 mm^2^, which was achieved by combining a 2.5× objective with optical zoom, allowing observation of the entire 6 mm diameter culture. The recording of spontaneous activity lasted for 15 minutes and was controlled using the Hokawo 2.10 software (Hamamatsu Photonics). During recordings, the cultured neuronal networks were placed in a glass microincubator (Ibidi GmbH), which maintained the same environmental conditions as the standard incubator. The temperature was set to 25°C to facilitate spontaneous activity.

### Induction of damage in the neuronal cultures

The local injury was induced by partially cutting a population of neurons on the 6 mm diameter PDMS using a scalpel on DIV 12 or 13. The protocol for monitoring network behavior is as follows: spontaneous activity was recorded for 15 minutes before injury. Immediately after injury, neuronal activity was recorded for 30 minutes. Activity was recorded for 15 minutes at quasi-logarithmic time points post-injury, specifically at 2, 6, and 24 h (Ayasreh et al., 2022).

### Activity quantification

On the images of acquired recording, a series of 150 × 150 μm^2^ square regions of calcium intensity data were defined as regions of interest (ROIs), resulting in a total of 1,400 regions. Spike trains of neuronal activity were extracted from each region by first generating a time series of mean fluorescence intensity within each ROI and then detecting inferred spiking events using a Schmitt-trigger filter (lower threshold = 0.4 and upper threshold = 0.8) (Soriano, 2023).

Neuronal activity comprised either individual activation of ROIs or coordinated episodes in which several ROIs displayed activity together within a short time window. These coordinated episodes were termed *network bursts*, which reflect the synchronized phenomenon of high-frequency firing occurring over a short period, representing the coordinated and rhythmic activity of neuronal populations (Wagenaar et al., 2006; Yamamoto et al., 2016; Pasquale et al., 2017). In our study, network bursts were identified when the number of spikes from the neuronal population within a 200 ms window exceeded 20% of the total ROIs, as below this fraction, a burst could not be easily distinguished from random background activity. The number of bursts during a recording and their amplitude (network fraction) were used to quantify neuronal culture activity before and after injury, as well as during recovery.

### Effective connectivity

Causal interactions between pairs of active ROIs were inferred using generalized transfer entropy (Stetter et al., 2012; Tibau et al., 2020; Montalà-Flaquer et al., 2022). Given the spike trains of a pair of ROIs *X* and *Y*, with *X* = {*x*_n_} and *Y* = {*y*_n_}, the amount of information transferred from ROI *Y* to *X* is given by:

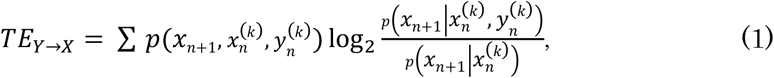

where *n* is a discrete-time index and *k* (= 2) is the Markov order. For inference, the 30-minute-long spike train of each ROI was built using a time bin of 20 ms, with data containing either ‘1’ for the presence of a spike or ‘0’ for its absence. Transfer entropy is a nonlinear and nonsymmetric measure in *X* and *Y* (TE_Y → X_ ≠ TE_X → Y_), allowing us to estimate directed interactions, i.e., effective connectivity, in the network. Significance was established by first normalizing the distribution of the TE values using a *z-score* transformation:

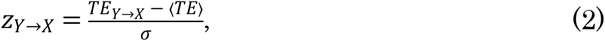

where ⟨*TE*⟩ is the mean value of all TE scores and *σ* their standard deviation, and then by setting a threshold *z*_*th*_ = 2 so that *z*_*Y*→*X*_ = 1 ∀ *z*_*Y*→*X*_ ≥ *z*_*th*_, and *z*_*Y*→*X*_ = 0 otherwise. The final adjacency matrix of the effective connections was directed and binary.

### *In silico* neuron model

The Izhikevich model was used in this study (Izhikevich, 2003) since it accurately reproduces the spiking behavior of cultured neurons with sufficient biological fidelity (Orlandi et al., 2013). The model is described by the following equation:

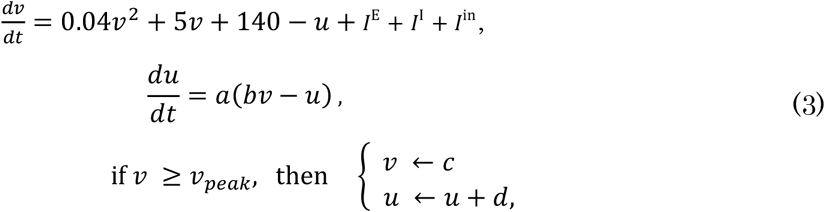

where, *v* is the membrane potential, *u* is the recovery variable, *I*^E^ is the excitatory synaptic input, *I*^I^ is the inhibitory synaptic input, and *I*^in^ the external current. During spontaneous activity, an external current was absent, indicating that *I*^in^ = 0. The membrane potential was reset when it exceeded *ν*_*peak*_ = 30 mV, at which point an action potential (a spike) was recorded for that neuron. The parameters *a, b, c*, and *d* determine the dynamic characteristics of the modeled neurons. In the present study, the simulated neuronal networks contained 80% excitatory and 20% inhibitory neurons. For excitatory neurons, the parameters were set as [*a, b, c, d*] = [0.02, 0.2, −65.0, 8.0], whereas for inhibitory neurons they were set as [*a, b, c, d*] = [0.1, 0.2, −65.0, 2.0].

### *In silico* network generation and infliction of damage

Neurons were randomly positioned in a homogeneous manner within a circular area of 3 mm in diameter, with a density of 400 neurons/mm^2^, resulting in approximately 2,800 neurons. Following the method of Orlandi et al. (Orlandi et al., 2013), dendrites were modeled as circles centered on each neuronal soma with a radius of 150 μm, while axons were described through a growth process as follows. For each axon, its maximum length *𝓁* was first determined by sampling from a Rayleigh distribution with an average length of ⟨*𝓁*⟩ = 1.1 mm. Then, segments of 0.1 mm in length were placed in a concatenated manner along a pseudo-straight path, such that, at each growing step, a new axon segment slightly deviated from the direction of the previous one by an angle *θ*_i_, according to the following probability:

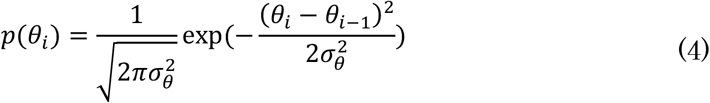

where *i* represents each growing step, and σ_θ_ was set to 5.73° (0.1 rad). The effects of PDMS on the development of the neuronal culture are modeled following (Houben et al., 2025). To account for the PDMS modulations, present in the living neuronal networks, virtual parallel bands 0.2 mm wide were considered for the “crevices” of the PDMS, which were separated by 0.3 mm wide “‘valleys.” These bands acted as obstacles that interfered with the axon growth, such that the axons crossed from top to bottom with a 50% probability and from bottom to top with a 5% probability. Whenever the axon failed to cross, it was deflected at an angle of 30° from the band. Simulations also included “control” scenarios with no virtual bands to investigate the impact of spatial constraints on network dynamics.

In both the “control” and “tracks” scenarios, a connection between neurons *i* → *j* was established whenever the axon of neuron *i* crossed the area covered by the dendritic tree of neuron *j* with a 20% probability. The resulting binary structural connectivity matrix ***A*** = {*a*_*ij*_} (1 for the presence of a connection and 0 otherwise) was then used to generate a new matrix ***W*** = {*w*_*ij*_} that preserved the connectivity relationships but incorporated weights. The specific values of *w*_*ij*_ are described in Section 2.10.

To model damage in the *in silico* networks, a straight line was defined as a reference in the center of the simulated culture, placed horizontally with a length of 1.5 mm (half the diameter of the network). Subsequently, all connections crossing this line were set to 0 in the connectivity matrix ***A***. Additionally, neurons whose axons were severed due to damage were considered dead (to mimic axonal transection) and excluded from further membrane potential updates during the simulation of the dynamics.

### Synaptic model

The numerical simulations utilized the following synaptic model, which is characterized by an exponential decay of induced postsynaptic membrane currents.

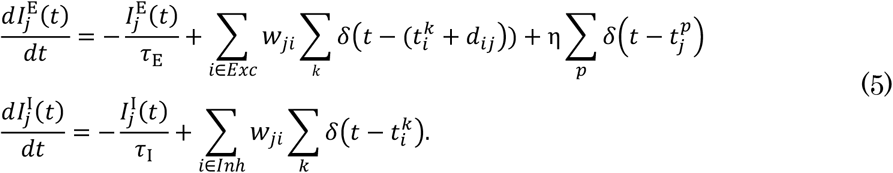

In this model, *j* is the postsynaptic neuron index, *t*_i_^*k*^ the time of the *k*-th somatic spike of presynaptic neuron *i*, and δ(·) the Dirac delta function. The set *Exc* refers to the excitatory neurons, and *Inh* refers to the inhibitory neurons. The parameters τ_E_ and τ_I_ account for the time constants of current loss and were set to τ_E_ = 5 ms for excitatory synapses and τ_I_ = 20 ms for inhibitory ones. Additionally, axonal conduction delays *d*_ij_ on excitatory connections were uniformly distributed in the range of [0, 5] ms, while those on inhibitory connections were fixed to 1 ms. The total excitatory and inhibitory input onto a neuron, *I*^E^ and *I*^I^, respectively, was the sum of synaptic currents from incoming input connections. Additionally, the target neuron *j* receives Poisson noise in the form of spontaneous synaptic inputs at an average rate of 1 Hz. The quantity *t*_j_^*p*^ denotes the time of the *p*-th Poisson spike input to neuron *j*, and η is the noise amplitude and was set to 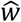, discussed in the next section.

### Spike-timing-dependent plasticity (STDP)

The STDP model (Rossum et al., 2000; Rubin et al., 2001; Gütig et al., 2003) is based on a phenomenon in which the strength of synaptic weight changes depending on the time difference between the firing of pre-synaptic and post-synaptic neurons. This is described mathematically by the following set of equations:

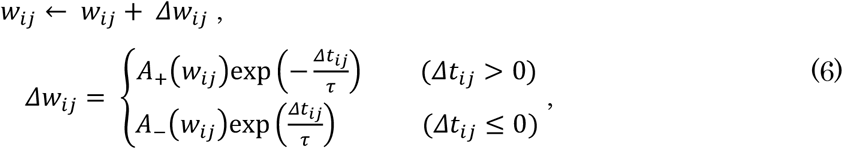

where *w*_*ij*_ is the synaptic weight from neuron *i* to *j* and *Δt* = *t*_*j*_ − *t*_*i*_ − *d*_*ij*_ the time difference between the firing of pre-synaptic neuron *i* and post-synaptic neuron *j. τ* is a constant set to 20 ms that accounts for the characteristic time for the strengthening or weakening of a connection. *A*_+_(*w*_*ij*_) and *A*_−_(*w*_*ij*_) are functions of *w*_*ij*_ (Rossum et al., 2000; Rubin et al., 2001; Gütig et al., 2003) given by:

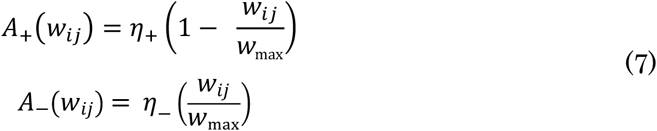

where *η*_+_ and *η*_-_ are learning constants, set to 0.1 and −0.12, respectively. *w*_max_ is the maximum value of the synaptic weight and is set to 6.8 unless otherwise specified. With *A*_+_(*w*_*ij*_) and *A*_−_(*w*_*ij*_) set in this way, the average value of synaptic weights is approximately 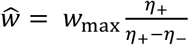. Since only the connection strengths between excitatory neurons were updated, all other connections were set to: 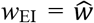 for excitatory-to-inhibitory connectivity, 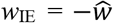 for inhibitory-to-excitatory, and 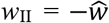 and for inhibitory-to-inhibitory. Nearest-neighbor spikes only contributed to modifying weights on plastic connections (Izhikevich and Desai, 2003; Morrison et al., 2008).

### Synaptic weight analysis

Network-wide connectivity through the synaptic weights matrix before and after injury was quantified by the global efficiency *E*_glob_ (Latora and Marchiori, 2001; Onnela et al., 2005; Fagiolo, 2007; Rubinov and Sporns, 2010) as:

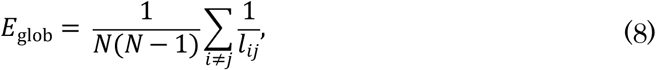

where *N* is the number of nodes in ***W***, and *l*_ij_ is the shortest path length between nodes *i* and *j*, obtained from path lengths *L*_ij_ through Dijkstra’s algorithm. *L*_ij_ was derived for the weighted connectivity matrix ***W*** as

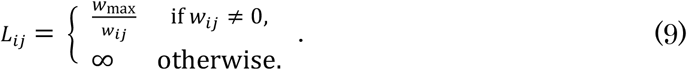

This procedure transforms weights into connection lengths with the property that higher weights correspond to shorter lengths.

### Normalized mutual information (NMI)

NMI is a measure of the similarity between two partitions (Alotaibi and Rhouma, 2022) and was used to quantify the differences in the community structure of the synaptic matrix ***W*** before and after damage:

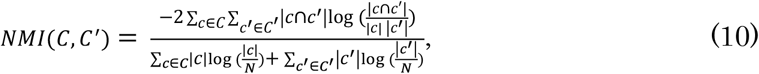

where *N* is the total number of neurons, and *C* is the best partition of ***W***, computed through the Louvain algorithm (Blondel et al., 2008). |*c*| and |*c’*| denote the number of neurons in the communities *c* and *c’* that belong, respectively, to partitions *C* and *C’*. |*c* ∩ *c′*| indicates the number of neurons in the intersection of *c* and *c′*. The greater the similarity in community partitions between two conditions, the higher the NMI.

### Reservoir Computing

The reservoir computing setup consisted of an input layer, a reservoir layer, and an output layer. Input signals were spoken digits from the TI-46 dataset (Liberman, Mark et al., 1993). The spoken digits were first converted to a 78-channel cochleagram using Lyon’s passive ear model (Slaney and Lyon, 1993). The cochleagram was then normalized to a maximum value of 1. The reservoir layer consists of a spiking neural network, with 5% of the total neurons receiving signals from the input layer. The signals delivered to each neuron were generated by summing two random channels of the 78-channel cochleagram. This input was then multiplied by the averaged synaptic weight *ŵ* and added to *I*^in^ in Eq. (3). During the task, the updating of synaptic weights was stopped. The reservoir state ***x***(*t*) was constructed from a subset of 5% of the neurons in the network, but different from the subset that received the inputs, according to the following equation:

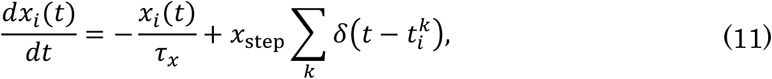

where *t*_i_^k^ is the time of the *k*-th spike of neuron *i*. The time constant *τ*_x_ was set to *τ*_x_ = 1 s, and *x*_step_ was set to 0.1. The output ***y***(*t*) was calculated as:

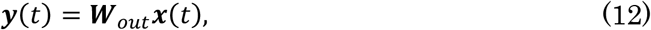

where ***W***_out_ is the output weight matrix, which was obtained using ridge regression during the training phase as:

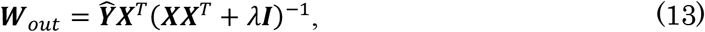

where ***X*** represents the reservoir state and was constructed by horizontally concatenating ***x***(*t*) from the start of the first trial to the end of the last trial. Each trial consisted of an input of a spoken digit, followed by a 10-second output period. The matrix 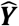 represents the target signal and was created by concatenating ***ŷ*** (*t*), a target vector, where the element corresponding to the correct label is set to 1 for 2.5 seconds following each stimulus onset and zero otherwise. The *λ* (= 1) is the regularization coefficient, and ***I*** is an identity matrix. The training dataset included 10 distinct samples of each spoken digit labeled as “zero,” “one,” and “two” from the speaker identified as “f1.” The test dataset was identical to the training dataset and was used to examine the loss and recovery of function due to damage. The estimated output was obtained from ***y***(*t*) as argmax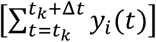, where *i* is an element of the output layer, *t*_k_ the onset of the *k*-th input, and *Δt* the target duration (set to 2.5 s). Accuracy was evaluated as the fraction of correct estimates.

## Results

### Self-organized recovery after injury in cultured and spiking neural networks

We constructed cultured neuronal networks with a modular structure by growing rat primary neurons on topographical PDMS substrates as described previously (Montalà-Flaquer et al., 2022) (Figure 1A). Crevices and valleys of the PDMS repeat in one direction, forming parallel tracks. There were approximately 10,000 neurons on a 6 mm diameter PDMS culture, and their behavior was monitored using calcium fluorescence imaging. The cultured neuronal networks were damaged at DIV 12 or 13 using a scalpel in a direction transverse to the PDMS tracks (Figure 1B). Subsequent recovery of neuronal activity was measured immediately after injury and at 15 min, 2 h, 6 h, and 24 h later. To mimic the experimental design, an SNN model was created to understand the synaptic and network mechanisms of the recovery process of the cultured neuronal network (Figure 1C). Network connectivity was modeled by simulating the growth process of nerve axons on a substrate that incorporated experimental-like topographical modulations. The created neural network models were initialized and then simulated with STDP for 72 simulated hours to settle all transients in the weight distribution. Then, the network models were damaged similarly to the culture experiments.

**Figure 1.**
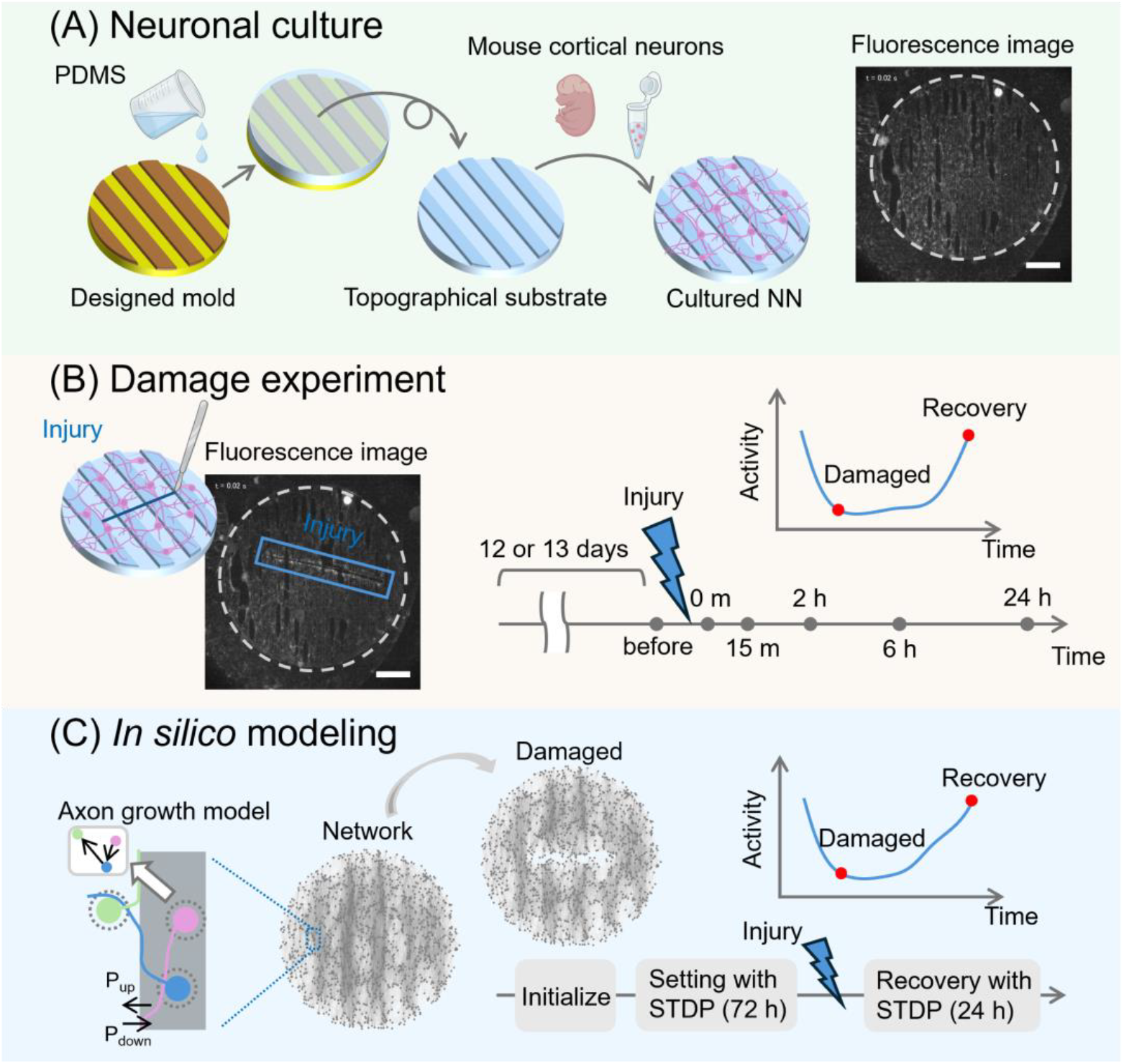
Schematic pipeline of the experimental and numerical approaches. **(A)** Primary cortical mouse neurons were cultured on a 6 mm diameter engineered substrate shaped with PDMS topographical modulations, and spontaneous neuronal activity was recorded using fluorescence calcium imaging. **(B)** Damage was applied to the culture with a scalpel, creating a wound approximately 4 mm long (blue box) at day *in vitro* 12-13. Spontaneous activity was then recorded just before the damage and at preset time points: immediately after the damage and at 15 minutes, 2 hours, 6 hours, and 24 hours post-injury, to monitor changes in activity levels and other properties. **(C)** In the *in silico* modeling, an axon growth model was used to replicate the structure of the cultured neuronal networks. Then, a spiking neural network model was initialized, and synaptic weights were adjusted using STDP over 72 hours. Injury was applied in the same manner as in the experiments, and network activity was evaluated by simulating STDP again at the same time points as in the cultured experiments. Scale bars in (A) and (B) are 1 mm.

Figure 2A shows fluorescence images of representative cultured neuronal networks before and after damage. Neuronal activity for each recording was extracted from the average fluorescence intensity of 1,400 ROIs that covered the culture area (see *Supplementary* Figure S1). The cultured neuronal network exhibited spontaneous activity characterized by collective quasi-synchronous events (network bursts). Such a bursting behavior was observed both before and immediately after damage (Figure 2B); however, in the latter condition network burst frequency was reduced. Effective connections were computed from firing patterns using transfer entropy, and functional communities were evaluated. These communities revealed groups of neurons that interacted more strongly within their groups than with the rest of the network.

**Figure 2.**
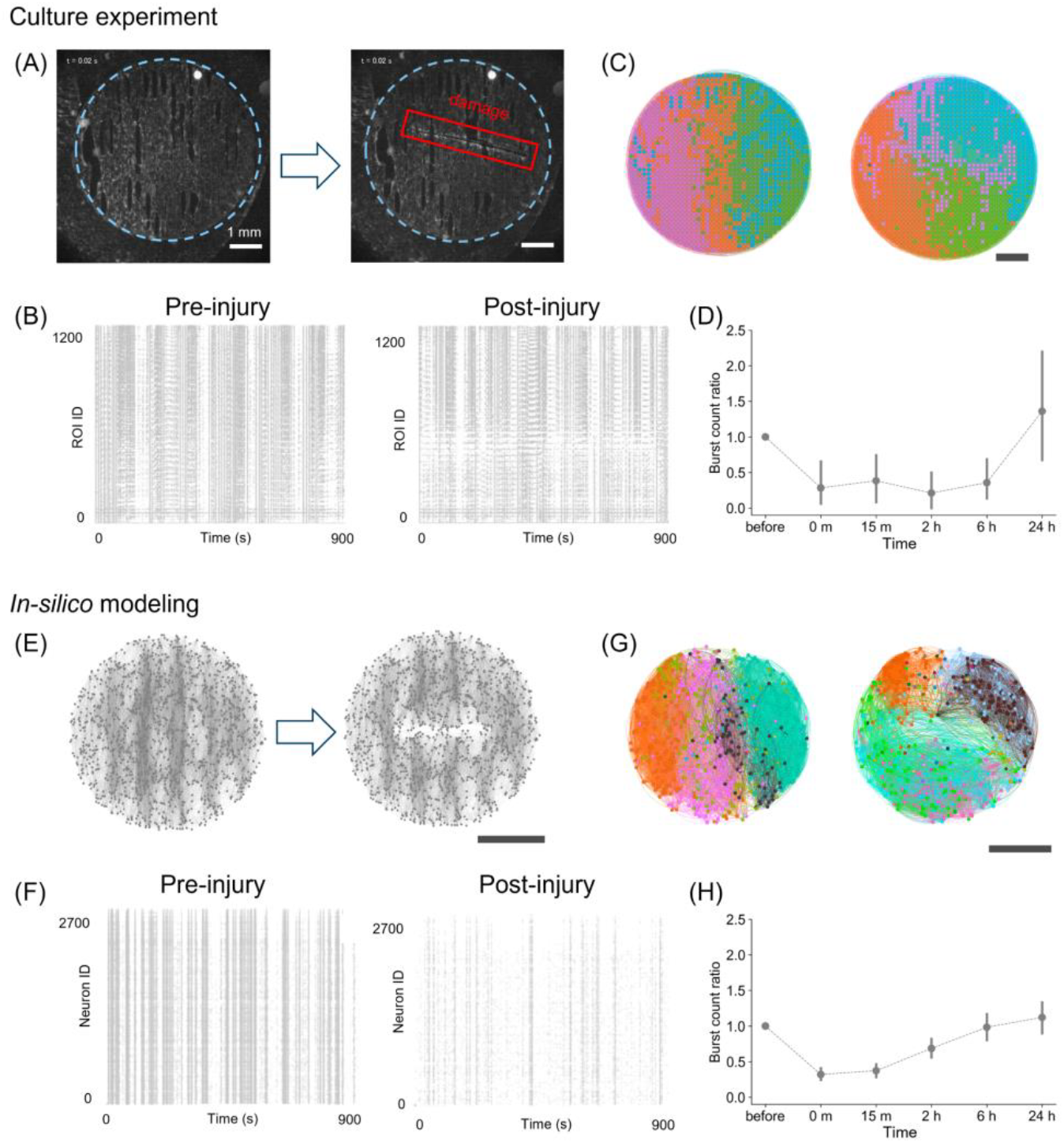
Damage and recovery in cultured neurons and *in silico*. **(A)** Fluorescence images of cultured neuronal networks before (left) and after damage (right). The blue dotted line highlights the PDMS engineered substrate where neurons grow, and the red box shows the area of damage. **(B)** Representative raster plots of neuronal activity in cultures before and immediately after injury. **(C)** Corresponding effective connectivity maps, illustrating that the functional communities tend to be aligned with the underlying PDMS topographical tracks before damage (left), but appear much less organized immediately after damage (right). **(D)** Ratio of bursting events after damage to before damage. The pre-injury state corresponds to a ratio of 1. After injury, collective activity abruptly decays to gradually recover afterwards. Error bars indicate the 95% confidence interval. **(E)** Illustrative network structural maps before and after injury in simulated neural networks. **(F)** Corresponding raster plots of neuronal activity before and immediately after damage in the simulations. **(G)** Effective connectivity maps and functional communities. As in the experiments, communities are initially aligned along tracks (left), but damage acutely causes them to become disorganized (right). **(H)** Ratio of bursting activity. A drop in activity followed by a post-damage recovery is observed, and very similar to the experimental observations. Scale bars in (A), (C), (E), and (G) are 1 mm.

As shown in Figure 2C, the spatial organization of the detected communities changed substantially throughout the network before and after injury (see *Supplementary* Figure S2). Before damage, the functional communities were aligned along the ‘tracks’ pattern, indicating that the substrate’s topographical modulation facilitated the formation of the communities. Such alignment was altered immediately after damage, suggesting that the local injury caused long-range effects. Interestingly, however, the cultured neurons reorganized their functional communities differently from the pre-damage communities, recovering the distribution of connection angles after 24 h (*Supplementary* Figure S2C).

Since activity in neuronal cultures is characterized by network bursts, changes in the frequency of such bursting were used as the first approach to quantify the damage. As shown in Figure 2D, the ‘burst count ratio,’ that is, the ratio of the number of bursting events relative to pre-damage conditions, substantially dropped by 0.72 immediately after injury. Upon recovery, the ratio approached pre-damage levels or even exceeded them 24 hours after injury.

To understand the synaptic mechanisms underlying this recovery, we utilized an SNN model, simulating a population consisting of 80% excitatory and 20% inhibitory neurons, where excitatory coupling was modulated by STDP (Figure 2E). Raster plots of activity were then generated, and the model parameters were adjusted —specifically the synaptic weights *w*_max_ — to qualitatively reproduce the experimental observations, e.g., a frequency of network bursts of 3.47 events/min (Figure 2F, left; see also *Supplementary* Figure S3), Finally, we conducted the same analysis pipeline as in the experiments to investigate whether the model captured the experimental observations on activity and effective connectivity. When damage was inflicted on the SNN model, a decrease in activity was observed (Figure 2F, right), and, as in the culture experiment, the organization of functional communities shifted from a track-oriented to an alternate arrangement immediately after damage (Figure 2G; *Supplementary* Figure S4) and returned to a track-oriented spatial structure after 24 hours (*Supplementary* Figure S4C). These results show that STDP in E-E synapses is sufficient to model the spontaneous activity and damage-induced changes in both dynamics and functional organization *in vitro*. Additionally, as in the culture experiments, quantification of the ratio of burst events showed a temporary decrease of 0.68 immediately after injury, with recovery to the original levels after approximately 24 hours of simulation (Figure 2H).

Functional recovery did not occur when STDP was absent (*Supplementary* Figure S5). However, we note that other plasticity mechanisms may also operate in this living system at once, e.g., synaptic scaling and homeostasis (Turrigiano, 1999, 2008; Effenberger et al., 2015).

### Reorganization of synaptic weights mediates recovery of neuronal activity *in silico*

To investigate the potential of STDP to spatially redistribute synaptic weights and restore activity, we analyzed the changes in excitatory-to-excitatory synaptic weights of the SNN model before and after injury. Figure 3A shows the change in synaptic weights of a representative network immediately after injury and 24 hours later. The increase and decrease in synaptic weights occurred throughout the entire network rather than at specific locations. We observed large fluctuations in synaptic weights post-damage, whereas the undamaged network remained relatively stable (Figure 3B), suggesting that injury enhanced the effect of STDP. Despite the large variation in the damaged network, the mean synaptic weight remained practically constant with a value of 3.08 before injury and after reorganization, indicating that network recovery did not require the formation of excessively strong weights (Figure. 3C).

**Figure 3.**
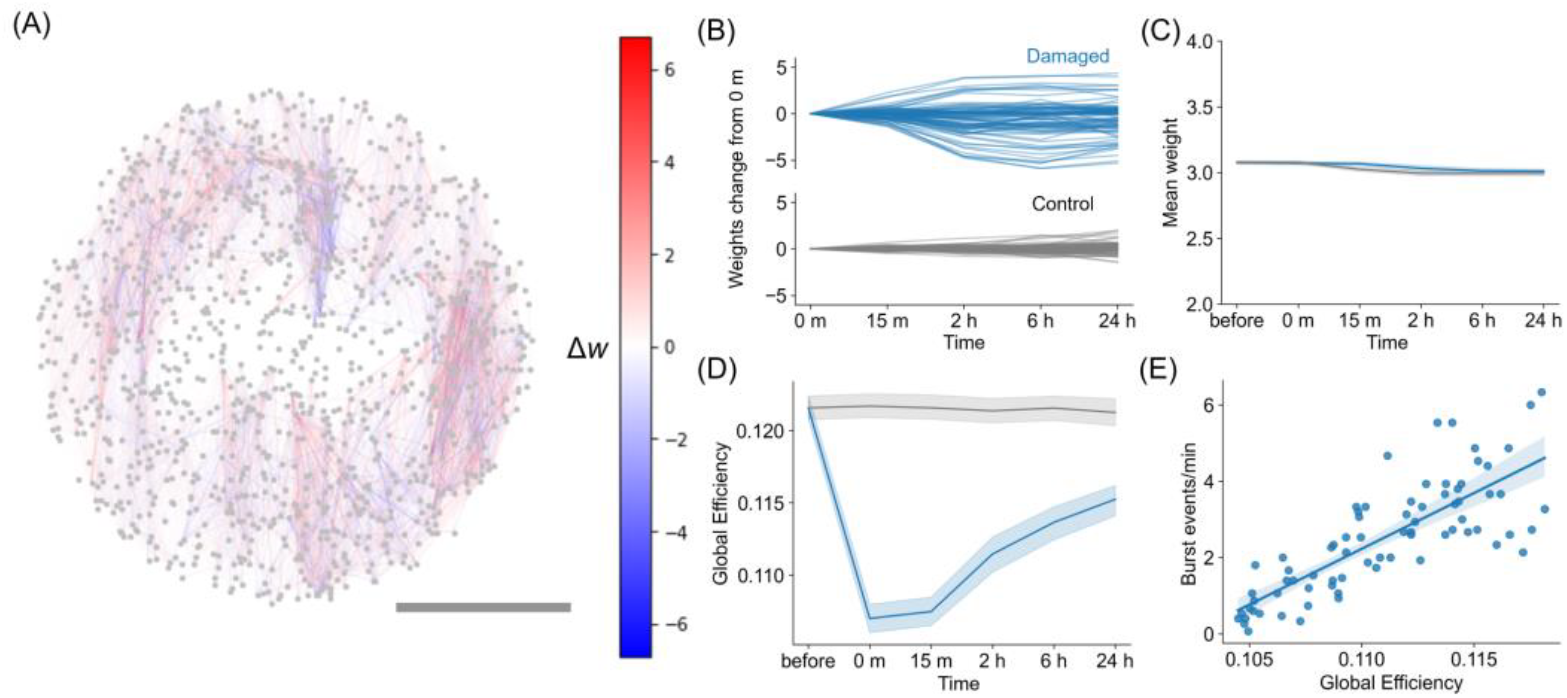
Changes in synaptic weights *in silico* due to STDP. **(A)** Representative network illustrating the changes in synaptic weights 24 h post-damage, as Δ*w* = w_24*h*_ − w_*injury*_. Red connections represent those that increase in weight, while blue connections represent those that decrease in weight. Scalebar is 1 mm. **(B)** Evolution of synaptic weights changes relative to the weights immediately after injury (0 min) for the representative damaged network and control network. The plot shows the 100 representative synaptic weights at each time point, with the values calculated by subtracting the weights at 0 min. **(C)** Evolution of the mean value of the synaptic weights for damaged networks and control networks, showing that the global change is minimal. The blue curve corresponds to a damaged network, whereas the grey corresponds to control network. Shadings indicate the 95% confidence interval, *n* = 15 networks. **(D)** Evolution of the global efficiency of the weighted connectivity matrix ***W*** after injury and during recovery. **(E)** Relationship between global efficiency and bursting event rate, with data exhibiting a Pearson’s correlation coefficient of *r* = 0.80 and *p* < 0.001 (two-sided Pearson correlation test, *n* = 75, degrees of freedom (*df*) = 73).

We then investigated the effect of post-damage reorganization of synaptic weights driven by STDP on the efficiency of information transfer in a network. We observed that the global efficiency of the damaged network decayed immediately after injury and gradually recovered over time (Figure 3D, blue curve), whereas the control condition exhibited only a small variation (grey curve). Global efficiency quantifies the fluency of neuronal communication across the entire network. Thus, the results indicate that communication between neurons was reduced due to damage but was restored through the reorganization of synaptic connections via STDP. Global efficiency was further examined in relation to the rate of network burst frequency, and we observed a strong correlation between the two quantities (*r* = 0.80, *p* < 0.001, two-sided Pearson correlation test, *n* = 75, degrees of freedom (*df*) = 73) (Figure. 3E). This correlation indicates that synchronous activity requires reliable communication between distant neurons in the network and, therefore, high network efficiency for information exchange. In summary, the STDP reorganization brings the post-damage network to a new state with increased global efficiency and activity level, but with different spatial distribution of connection weights.

### Effect of modular organization on damage and recovery in the model

We next examined the influence of the modular structure, shaped by a series of interconnected parallel tracks, to provide robustness against various damage scenarios. We designed several damage conditions, alongside control scenarios, in which either intra- or inter-modular connections were damaged, as shown in Figure 4A and *Supplementary* Figure S6. Cuts perpendicular to and along the tracks affected the intra-modular and inter-modular connections, respectively. A network generated by modeling axonal growth on an unpatterned surface was used as a control. Two different damage extents were considered: one in which the length of the injury extended half the diameter of the network and another in which the injury extended the full width (see Figure 4A).

**Figure 4.**
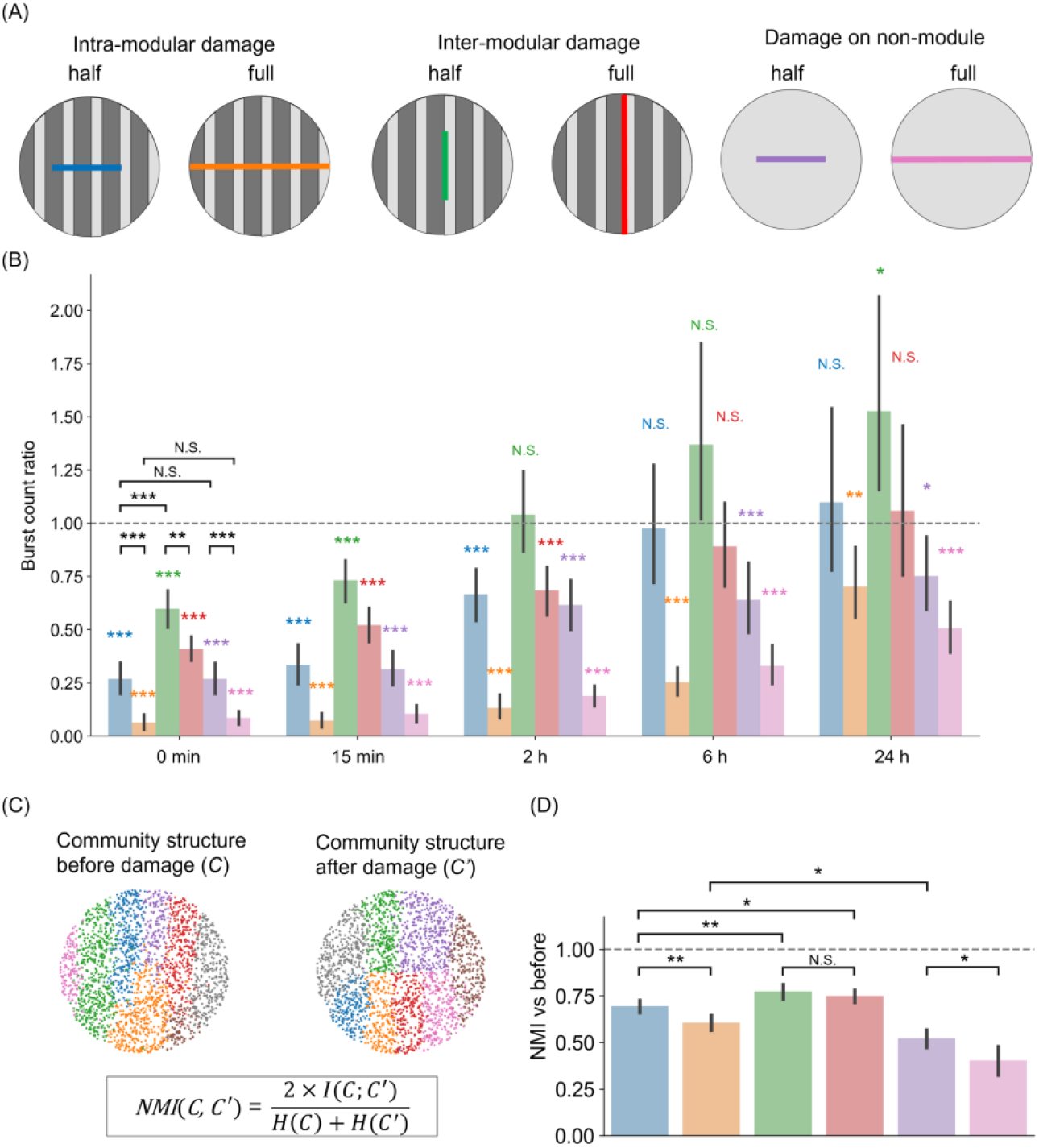
Importance of modularity on the magnitude and direction of damage in the model. **(A)** Schematic representation of damage actions on patterned (modular) and homogeneous networks. Intra-modular or inter-modular damages are achieved by, respectively, cutting the network along the direction perpendicular or parallel to the tracks’ orientation. In the sketch, ‘half’ indicates that the cut extends half the diameter of the network, while ‘full’ indicates that the damage effectually separates the network in two parts. ‘Damage on non-module’ represents the control scenario of neurons simulated on a flat surface. **(B)** Burst frequency changes for the six damage scenarios. For each scenario, the color bar shows the ratio of burst frequency between the damaged and control conditions. Color bars use the same color scheme as in (A). The bar height indicates the mean, and error bars indicate the 95% confidence intervals. Color asterisks indicate a two-sided one-sample *t*-test (^*^*p* < 0.05; ^**^*p* < 0.01; ^***^*p* < 0.001; N.S., no significance; *n* = 15, *df* = 14). Asterisks in black correspond to a two-sided unpaired *t*-test (^**^*p* < 0.01; ^***^*p* < 0.001; N.S., no significance; *n* = 15, *df* = 13). **(C)** Sketch illustrating the concept of normalized mutual information (NMI). Community structure is obtained via the Louvain algorithm. **(D)** The NMI values between the post- and pre-injury conditions for the six damage scenarios.

Figure 4B shows the rate of network burst relative to the non-damaged control at each time point after injury for a total of six conditions (four in modular networks and two in non-modular networks). Immediately after injury (0 min), the activity was lower than the control in all conditions (intra-modular half, 0.27 ± 0.14 (mean ± SD), two-sided one-sample *t*-test, *p* < 0.001 (vs. 1.00); intra-modular full, 0.06 ± 0.07, *p* < 0.001; inter-modular half, 0.60 ± 0.17, *p* < 0.001; inter-modular full, 0.41 ± 0.11, *p* < 0.001; non-module half, 0.27 ± 0.14, *p* < 0.001; non-module full, 0.09 ± 0.06, *p* < 0.001; *n* = 15, *df* = 14). A comparison between half-cuts and full-cuts showed that networks with a full-cut had significantly reduced activity (two-sided unpaired *t*-test, *p* < 0.001 (intra-modular half vs. intra-modular full); *p* < 0.01 (inter-modular half vs. inter-modular full); *p* < 0.001 (non-module half vs. non-module full)). Additionally, damage perpendicular to the tracks direction (intra-modular) had a greater impact on the rate of neuronal activity than damage applied along the tracks (inter-modular) (two-sided unpaired *t*-test, *p* < 0.001, intra-modular half vs. inter-modular half). However, there were no significant differences between non-modular networks and modular networks in which damage was applied within a module, i.e., perpendicular to the tracks orientation (two-sided unpaired *t*-test, *p* = 0.99 (intra-modular half vs. non-module half); *p* = 0.38 (intra-modular full vs. non-module full)). In summary, activity decreased as the size of the injury increased, and intra-modular damage had a greater effect than inter-modular damage.

The recoverability of the network was dependent on multiple factors, such as the size and direction of damage, and the presence of modular structure. When the injury was small and applied along the tracks (green bar in Figure 4B), activity recovered to the pre-damage level 2 h later (inter-modular half, 1.04 ± 0.38, two-sided one-sample *t*-test, *p* = 0.71 (vs. 1.00)). This recovery was further observed at 6 h, with activity comparable to that without damage in the cases of damage along tracks (green and red bars) and half damage across tracks (blue bar) (intra-modular half, 0.98 ± 0.57, *p* = 0.87; inter-modular full, 0.89 ± 0.40, *p* = 0.32). However, even after 24 h, the activity did not recover in the case of full intra-modular damage (orange bar), i.e., across tracks (intra-modular full, 0.70 ± 0.33, *p* < 0.05). Furthermore, in the absence of a specific modular structure (purple and pink bars), activity was not restored, regardless of damage size (non-module half, 0.75 ± 0.33, *p* < 0.05; non-module full, 0.51 ± 0.25, *p* < 0.001). The differences in damage and recovery caused by network structure and cut orientations were also observed when the same number of connections were eliminated (*Supplementary* Figure S7). These findings suggest that modular structures facilitate recovery from local injuries.

To examine differences in damage recovery depending on the presence or absence of a modular structure, we focused on community structure before and after damage. We quantified the differences in community structure using normalized mutual information (NMI) (Figure 4C). Immediately after injury, the NMI deviated from 1.0 under all the damaged conditions (Figure 4D), indicating that the community structure was altered by the injury. The size of the injury strongly affected the communities in the case of intra-modular damage and in the non-modular scenario (two-sided unpaired *t*-test, *p* < 0.01 (intra-modular half vs. intra-modular full); *p* < 0.05 (non-module half vs. non-module full), *n* = 15). However, cut size did not affect the communities in the case of inter-modular damage (two-sided unpaired *t*-test, *p* = 0.30 (inter-modular half vs. inter-modular full)). In the context of the tracks pattern, damage across tracks had a more significant impact than damage along the tracks (two-sided unpaired *t*-test, *p* < 0.01 (intra-modular half vs. inter-modular half); *p* < 0.05 (intra-modular half vs. inter-modular full)). Among these damages on modular networks, the strongest influence was in the networks with full intra-modular damage, but they had greater NMI than networks without a specific modular structure (two-sided unpaired *t*-test, *p* < 0.05 (intra-modular full vs. non-module half)). In summary, we conclude that modular structures reduce the alteration of community structures due to damage, leading to faster recovery than non-modular networks.

### Restoration of information representation and processing through the recovery of neural network activity

Finally, we employed a reservoir computing framework to explore the computational impact of network recovery using STDP (Figure 5A). In the reservoir computing framework used here, a signal is injected into the network, and its response is linearly decoded to perform a classification task. Spoken digits were used as sensory information and served as input to 5% of the neurons in the SNN both before and after damage. Figure 5B shows the neural responses to the spoken digit “zero” before and after full damage across tracks. Before the injury, the neurons responded well to the input, and the summed activity of the neurons produced two peaks. These peaks of summed neural activity aligned with the peaks of the summed spectrogram of the input spoken digits with a time delay. After injury, the neural response decreased, and the two peaks became smaller. The magnitude of the response recovered after 24 hours. This indicates that self-organization through synaptic plasticity restores not only spontaneous activity but also the capacity of the system to respond to external inputs.

**Figure 5.**
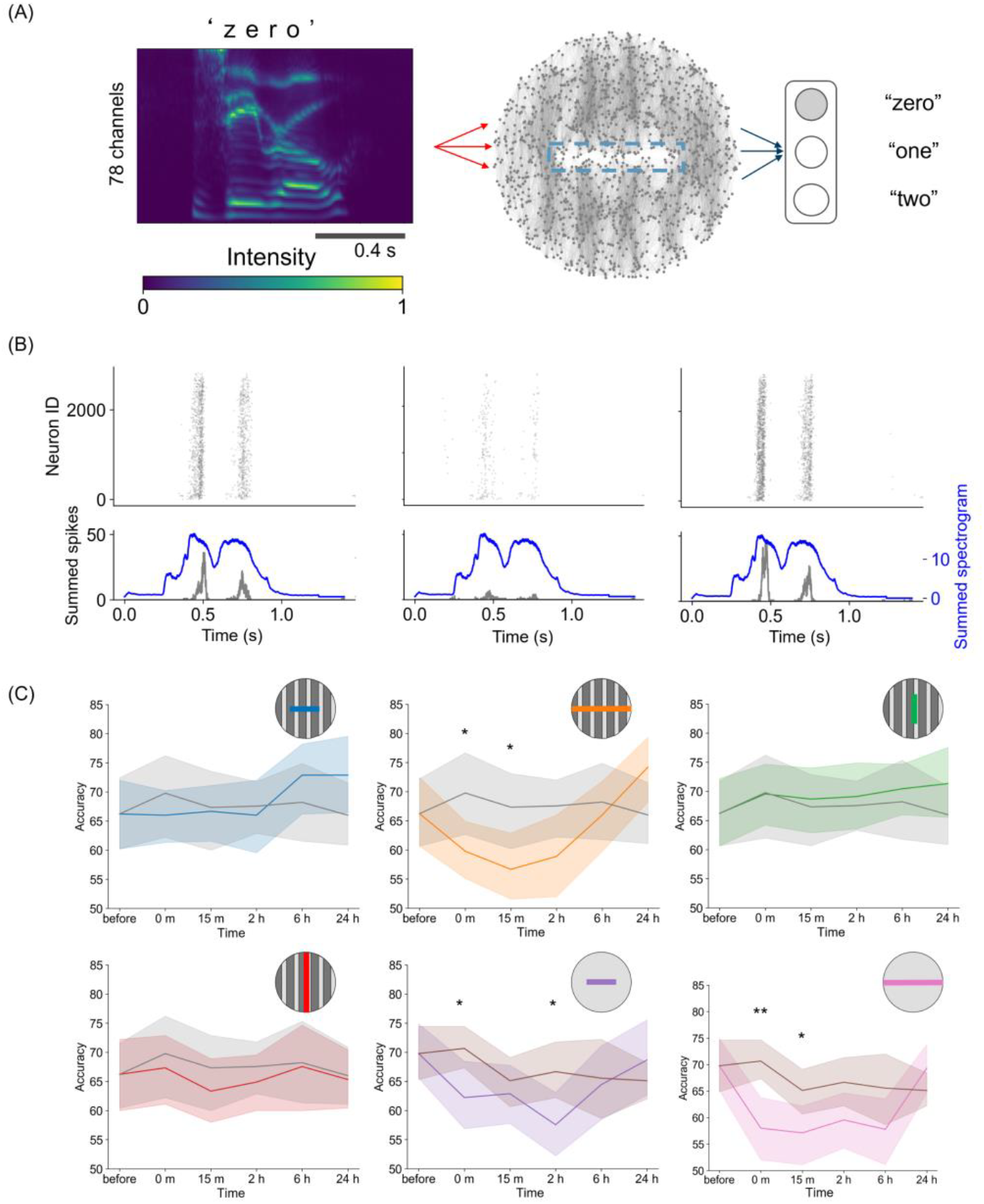
Reservoir computing on a damaged neural network. **(A)** Sketch of the reservoir computing framework. The spectrogram of spoken digits is delivered into a spiking neural network, and the evoked activity is regressed as output. **(B)** Raster plots of evoked activity for the spoken digit “zero” before (left), immediately after (center), and 24 hours after injury (right). The bottom panels show the summed-up spikes (black) and the spectrogram (blue). **(C)** Time courses of accuracy in the reservoir computing tasks for the six damage conditions. The colors correspond to each type of damage. Grey curves show the undamaged case for the tracks-patterned network, and the brown curves show the undamaged case for the control, unpatterned network. In the panels, the lines indicate the average value, and the shaded areas represent the 95% confidence interval. ^*^*p* < 0.05; ^**^*p* < 0.01; ^***^*p* < 0.001 (two-sided unpaired *t-*test, *n* = 15, *df* = 13).

Subsequently, we used multiple spoken digits to perform classification tasks. In this task, the linear decoder in the output layer was trained to obtain the optimal output weight matrix ***W***_out_ to classify the spoken digits “zero,” “one,” and “two.” When the output weight matrix was trained on the network responses before damage, the classification accuracies were 66.2 ± 11.6% (mean ± SD) for the tracks-patterned network and 69.8 ± 9.5% for the unpatterned network, both of which were significantly higher than a random guess (33.3%). When the output weights were fixed at the pre-damage state, classification accuracy decreased over time, regardless of the presence of damage (*Supplementary* Figure S8). This decline occurred because STDP alters the synaptic weights of the reservoir’s SNN, resulting in changes in the information representation.

Furthermore, we retrained the output weight matrix for each damaged and recovered network state. In the undamaged case, performance was maintained when ***W***_out_ was retrained (Figure 5C). This indicates that the input information was quantitatively preserved in the neural response and could be classified by a linear decoder, even though the information representation changed over time due to plasticity. Such maintenance of performance was observed under three of the four conditions applied to the modular neural network (intra-modular half, inter-modular half, and inter-modular full), suggesting that the modular structure is robust against small damage in terms of sensory information representation. In contrast, under the conditions of damage on non-module and full intra-modular damage, accuracy dropped immediately after damage (intra-modular full, 59.8 ± 10.1%, *p* < 0.05; non-module half, 62.2 ± 11.5%, *p* < 0.05; non-module full, 58.0 ± 11.2%, *p* < 0.01; *n* = 15). In other words, under these conditions, damage impaired the network’s ability to represent information and perform speech recognition as a reservoir. While such temporary dysfunction was observed, classification accuracy gradually recovered over time and returned to its original level after 24 h. This suggests that information representation in neural networks improves with the recovery of synchronous activity.

## Discussion

In the present study, we developed a cultured neuronal network with a modular structure using topographical substrates. Modularity is a distinctive feature of the brain, evolutionarily conserved across many species (Meunier et al., 2010), and is believed to be a crucial trait in living neuronal circuits (Sporns and Betzel, 2016; Michaels et al., 2020; Vishwanathan et al., 2024). Recent studies in neuronal network patterning *in vitro* have shown that developing a modular structure in neuronal networks can generate rich activity that mimics the complex information processing of the brain (Yamamoto et al., 2018, 2023; Montalà-Flaquer et al., 2022). PDMS plays a critical role in neuronal culture. Microfluidic devices made with PDMS can control the structure of neuronal networks (Takemuro et al., 2020), and culturing cortical neurons on PDMS can reduce mechanical mismatch (Sumi et al., 2020) and adjust the topology of the network through topographical processing of PDMS (Montalà-Flaquer et al., 2022). Furthermore, when damaging neuronal networks, PDMS provides a scaffold that allows the mechanical damage to be applied with a scalpel (Ayasreh et al., 2022).

In this study, we first examined the changes in activity caused by damage to cultured neuronal networks, extending previous studies that also investigated the impact of damage. These studies used laser microdissection or a scalpel to damage a subpopulation of the network (Teller et al., 2020; Ayasreh et al., 2022), while others focused on the modulation of structural connectivity through heat (Hong and Nam, 2020). In these studies, the authors observed that cultured neuronal networks temporarily decreased the number of activations and the rate of synchronous activity when damaged, although activity was restored within minutes or days, depending on the magnitude of the damage. However, the overall recovery was not homogeneous; the area of the network surrounding the damage recovered well, whereas the damaged area remained silent and was effectively unrecoverable. Our study showed that immediately after injury, the rate of activity in the network decreased but recovered after 24 hours (Figure 2D), which is consistent with previous studies.

Motivated by these culture experiments, we extended our study using a spiking neural network model. The combination of mathematical models and culture experiments facilitated the design of extended *in silico* experiments, allowing us to compare a wide range of conditions. Specifically, numerical simulations enabled us to explore the mechanisms underlying the restoration of neuronal network activity, which may involve the reorganization of the neuronal network through plasticity. STDP is a natural candidate for such plasticity because it plays important remodeling roles in various regions of the brain (Bi and Poo, 1998; Schmidgall et al., 2024) and has led to the restoration of activity in simulations of randomly inactivating neurons (Gabrieli et al., 2020). Even in cases of severe network disconnection, as in our experiments, STDP worked well and was sufficient to restore neuronal activity (Figure 2H). Further investigation of synaptic weights revealed that synaptic plasticity enhanced global efficiency in the damaged network (Figure 3D). Although previous studies using cultured neuronal networks have estimated the recovery of global efficiency by inferring effective connectivity from network activity (Teller et al., 2020; Ayasreh et al., 2022), our studies have shown that it is also possible to examine the synaptic weights underlying firing patterns using a mathematical model that replicates cultured neuronal networks. This agreement provides evidence that our model reproduces the culture experiments well and opens new avenues for monitoring unknown parameters, such as synaptic weights, which cannot be fully examined in culture experiments.

The proposed model and its capacity to successfully reproduce the experimental observations allowed us to extend the simulations to explore the effects of different damage conditions. The results showed that the effect of damage in intra-modular conditions was stronger than in inter-modular conditions and that networks without modular organization were unable to fully recover their activity (Figure 4B). In this regard, we suggest that recovery can be achieved by reorganizing new functional communities that by regions activating together. Although the new communities departed from the original ones, the underlying initial modular configuration seemed to provide a backbone to facilitate the maintenance of functional communities (Figure 4D). This hypothesis is supported by the fact that modular organization is known to be a damage-resistant structure due to its functional separation and redundancy (Zhang et al., 2019; Chen et al., 2021). For instance, animal experiments have shown that rodents can efficiently perform short-term memory tasks when artificially perturbed by optogenetic stimulation in the brain hemisphere; however, a modular organization is required for the robustness of persistent activity in response to perturbations (Chen et al., 2021). Additionally, in the present study, the neuronal activity of some samples, particularly in the half inter-modular damage cases, was significantly greater than the control level 24 hours after injury. (Figure 4B). Such overactivity may be associated with clinical studies showing that epilepsy can occur after injury (Herman, 2002; Englander et al., 2014; Ding et al., 2016). Further time-tracking is expected to enable the investigation of chronic mechanisms.

Finally, we observed an association between the restoration of synchronous activity and the recovery of cognitive-like functions, such as spoken digits classification, using a reservoir computing framework. Reservoir computing can be integrated with cultured or computational neural networks to link neuronal dynamics to cognitive and behavioral tasks (Nicola and Clopath, 2017; Yada et al., 2021; Cai et al., 2023; Sumi et al., 2023). In the present study, we used this framework to investigate how changes in neuronal dynamics before and after injury affect speech signal classification. Our model demonstrated that the modular structure was functionally robust to relatively small injuries and that functional impairment, even due to serious damage, could be restored if down-stream neuronal circuits are able to re-organize as well (Figure 5C). Although injury-induced cognitive impairment and recovery have been observed in animal studies (Chen et al., 2010; Christensen et al., 2008; Edlow et al., 2021), our results provide a new perspective on how the dynamic process of synaptic reorganization after injury affects information representation and processing at the neuronal network level. The reservoir computing framework can also be employed for other tasks, such as motor learning and memory, by adjusting the readout. Therefore, it can be applicable to model various neurological dysfunctions and recovery. In our study, functional recovery was achieved through spontaneous synaptic reorganization of functional connectivity and task-dependent learning in the output layer. However, in clinical rehabilitation, patients modify the functional connectivity of the motor cortex through input-output learning in addition to spontaneous modulation (Murata et al., 2015). The recovery through input-output interactions is reported to be promoted by repeated 40 Hz stimulation, which rescues synaptic plasticity such as STDP (Wang et al., 2023). Also, a theoretical study has shown that repeated stimulation inputs to the STDP model can lead to the formation of temporal patterns (Hosaka et al., 2008). The ongoing development of these fields is expected to lead to future applications in rehabilitation models using repetitive input and synaptic plasticity in the neuronal network to restore function.

To conclude, it is worth emphasizing that damage to neuronal networks occurs in several pathologies (Carmichael, 2016; Nagappan et al., 2020). In our study, a cultured neuronal network was designed as an accessible laboratory model for damage to neuronal circuits, which was successfully reproduced computationally. Our model has the potential to predict changes in functional neuronal networks and dynamics due to local damage and may be useful in designing advanced models that combine spontaneous and evoked activity to treat dysfunction. In addition, the mathematical description of the self-organized recovery process can be applied to the field of information processing. The brain is often compared to an electronic computer as an information-processing device. However, damage resistance and recovery capabilities are unique characteristics of living neuronal networks, particularly in the brain (Hassabis et al., 2017). Analysis of recovery from damage through cultured neuronal networks and mathematical neural models is valuable for understanding the mechanisms of neuronal function recovery and for designing artificial neural networks with damage tolerance and self-repair capabilities (Psaier and Dustdar, 2011; Khlaisamniang et al., 2023). Our findings could contribute to further research to introduce self-repair capabilities in robots and AI systems and to advance the understanding of diseases caused by local injury and their treatments in the human nervous system.

## Acknowledgment

This work was supported by the MEXT Grant-in-Aid for Transformative Research Areas (A) and (B) “Multicellular Neurobiocomputing” (21H05163, 21H05164, 24H02330, 24H02332, 24H02334), JSPS KAKENHI (21K12050, 22KK0177, 23H00251, 22KJ0190, 23K28179), JST Moonshot R&D Program (JPMJMS2023), JSPS Overseas Challenge Program for Young Researchers, and the Tohoku University Research Institute of Electrical Communication (RIEC) Cooperative Research Project Program. This research was supported by the European Union Horizon 2020 research and innovation program under Grant No. 964877 (project NEU-CHiP). J.S. acknowledges support from grant PID2022-137713NB-C22, funded by MCIU/AEI/10.13039/501100011033, ERDF/EU, and the Generalitat de Catalunya under grant 2021-SGR-00450. Cartoons in Figure 1A and B were created with BioRender.com.

## Author Contributions

T.S., H.Y., and J.S. designed the experiments; T.S. and A.M.H. performed the experiments; T.S., A.M.H., and J.S. analyzed the data; H.K. and Y.K. contributed new analytical tools; T.S., H.Y., and A.H.-I. wrote the initial draft of the manuscript.

## Notes

### Competing Interest Statement

The authors have declared no competing interest.

## Reference

Alotaibi, N., and Rhouma, D. (2022). A review on community structures detection in time evolving social networks. Journal of King Saud University - Computer and Information Sciences 34, 5646–5662. doi: 10.1016/j.jksuci.2021.08.016

Arnemann, K. L., Chen, A. J.-W., Novakovic-Agopian, T., Gratton, C., Nomura, E. M., and D’Esposito, M. (2015). Functional brain network modularity predicts response to cognitive training after brain injury. Neurology 84, 1568–1574. doi: 10.1212/WNL.0000000000001476

Ayasreh, S., Jurado, I., López-León, C. F., Montalà-Flaquer, M., and Soriano, J. (2022). Dynamic and Functional Alterations of Neuronal Networks In Vitro upon Physical Damage: A Proof of Concept. Micromachines 13, 2259. doi: 10.3390/mi13122259

Bi, G., and Poo, M. (1998). Synaptic Modifications in Cultured Hippocampal Neurons: Dependence on Spike Timing, Synaptic Strength, and Postsynaptic Cell Type. J. Neurosci. 18, 10464–10472. doi: 10.1523/JNEUROSCI.18-24-10464.1998

Blondel, V. D., Guillaume, J.-L., Lambiotte, R., and Lefebvre, E. (2008). Fast unfolding of communities in large networks. J. Stat. Mech. 2008, P10008. doi: 10.1088/1742-5468/2008/10/P10008

Boroda, E., Armstrong, M., Gilmore, C. S., Gentz, C., Fenske, A., Fiecas, M., et al. (2021). Network topology changes in chronic mild traumatic brain injury (mTBI). NeuroImage: Clinical 31, 102691. doi: 10.1016/j.nicl.2021.102691

Bunday, K. L., and Perez, M. A. (2012). Motor Recovery after Spinal Cord Injury Enhanced by Strengthening Corticospinal Synaptic Transmission. Curr Biol 22, 2355–2361. doi: 10.1016/j.cub.2012.10.046

Cai, H., Ao, Z., Tian, C., Wu, Z., Liu, H., Tchieu, J., et al. (2023). Brain organoid reservoir computing for artificial intelligence. Nat Electron 6, 1032–1039. doi: 10.1038/s41928-023-01069-w

Carmichael, S. T. (2016). The 3 Rs of Stroke Biology: Radial, Relayed, and Regenerative. Neurotherapeutics 13, 348–359. doi: 10.1007/s13311-015-0408-0

Chen, G., Kang, B., Lindsey, J., Druckmann, S., and Li, N. (2021). Modularity and robustness of frontal cortical networks. Cell 184, 3717-3730.e24. doi: 10.1016/j.cell.2021.05.026

Chen, H., Epstein, J., and Stern, E. (2010). Neural Plasticity After Acquired Brain Injury: Evidence from Functional Neuroimaging. PM&R 2, S306–S312. doi: 10.1016/j.pmrj.2010.10.006

Christensen, B. K., Colella, B., Inness, E., Hebert, D., Monette, G., Bayley, M., et al. (2008). Recovery of Cognitive Function After Traumatic Brain Injury: A Multilevel Modeling Analysis of Canadian Outcomes. Archives of Physical Medicine and Rehabilitation 89, S3–S15. doi: 10.1016/j.apmr.2008.10.002

Ding, K., Gupta, P. K., and Diaz-Arrastia, R. (2016). “Epilepsy after Traumatic Brain Injury,” in Translational Research in Traumatic Brain Injury, eds. D. Laskowitz and G. Grant (Boca Raton (FL): CRC Press/Taylor and Francis Group). Available at: http://www.ncbi.nlm.nih.gov/books/NBK326716/ (Accessed August 27, 2024).

Edlow, B. L., Claassen, J., Schiff, N. D., and Greer, D. M. (2021). Recovery from disorders of consciousness: mechanisms, prognosis and emerging therapies. Nat Rev Neurol 17, 135–156. doi: 10.1038/s41582-020-00428-x

Effenberger, F., Jost, J., and Levina, A. (2015). Self-organization in Balanced State Networks by STDP and Homeostatic Plasticity. PLOS Computational Biology 11, e1004420. doi: 10.1371/journal.pcbi.1004420

Englander, J., Cifu, D. X., and Diaz-Arrastia, R. (2014). Seizures after Traumatic Brain Injury. Arch Phys Med Rehabil 95, 1223–1224. doi: 10.1016/j.apmr.2013.06.002

Fagiolo, G. (2007). Clustering in complex directed networks. Phys. Rev. E 76, 026107. doi: 10.1103/PhysRevE.76.026107

Fukui, A., Osaki, H., Ueta, Y., Kobayashi, K., Muragaki, Y., Kawamata, T., et al. (2020). Layer-specific sensory processing impairment in the primary somatosensory cortex after motor cortex infarction. Sci Rep 10, 3771. doi: 10.1038/s41598-020-60662-7

Gabrieli, D., Schumm, S. N., Vigilante, N. F., Parvesse, B., and Meaney, D. F. (2020). Neurodegeneration exposes firing rate dependent effects on oscillation dynamics in computational neural networks. PLoS One 15, e0234749. doi: 10.1371/journal.pone.0234749

Gütig, R., Aharonov, R., Rotter, S., and Sompolinsky, H. (2003). Learning Input Correlations through Nonlinear Temporally Asymmetric Hebbian Plasticity. J. Neurosci. 23, 3697–3714. doi: 10.1523/JNEUROSCI.23-09-03697.2003

Hassabis, D., Kumaran, D., Summerfield, C., and Botvinick, M. (2017). Neuroscience-Inspired Artificial Intelligence. Neuron 95, 245–258. doi: 10.1016/j.neuron.2017.06.011

Herman, S. T. (2002). Epilepsy after brain insult. Neurology 59, S21–S26. doi: 10.1212/WNL.59.9_suppl_5.S21

Hong, N., and Nam, Y. (2020). Thermoplasmonic neural chip platform for in situ manipulation of neuronal connections in vitro. Nat Commun 11, 6313. doi: 10.1038/s41467-020-20060-z

Hosaka, R., Araki, O., and Ikeguchi, T. (2008). STDP provides the substrate for igniting synfire chains by spatiotemporal input patterns. Neural Comput 20, 415–435. doi: 10.1162/neco.2007.11-05-043

Houben, A. M., Garcia-Ojalvo, J., and Soriano, J. (2025). Role of connectivity anisotropies in the dynamics of cultured neuronal networks. doi: 10.48550/arXiv.2501.04427

Izhikevich, E. M. (2003). Simple model of spiking neurons. IEEE Transactions on Neural Networks 14, 1569–1572. doi: 10.1109/TNN.2003.820440

Izhikevich, E. M., and Desai, N. S. (2003). Relating STDP to BCM. Neural Computation 15, 1511–1523. doi: 10.1162/089976603321891783

Khlaisamniang, P., Khomduean, P., Saetan, K., and Wonglapsuwan, S. (2023). Generative AI for Self-Healing Systems., in 2023 18th International Joint Symposium on Artificial Intelligence and Natural Language Processing (iSAI-NLP), 1–6. doi: 10.1109/iSAI-NLP60301.2023.10354608

Latora, V., and Marchiori, M. (2001). Efficient Behavior of Small-World Networks. Phys. Rev. Lett. 87, 198701. doi: 10.1103/PhysRevLett.87.198701

Lee, W.-C. A., Bonin, V., Reed, M., Graham, B. J., Hood, G., Glattfelder, K., et al. (2016). Anatomy and function of an excitatory network in the visual cortex. Nature 532, 370–374. doi: 10.1038/nature17192

Liberman, Mark, Amsler, Robert, Church, Ken, Fox, Ed, Hafner, Carole, Klavans, Judy, et al. (1993). TI 46-Word. doi: 10.35111/ZX7A-FW03

Lynn, C. W., and Bassett, D. S. (2019). The physics of brain network structure, function and control. Nat Rev Phys 1, 318–332. doi: 10.1038/s42254-019-0040-8

Meunier, D., Lambiotte, R., and Bullmore, E. T. (2010). Modular and Hierarchically Modular Organization of Brain Networks. Front. Neurosci. 4. doi: 10.3389/fnins.2010.00200

Michaels, J. A., Schaffelhofer, S., Agudelo-Toro, A., and Scherberger, H. (2020). A goal-driven modular neural network predicts parietofrontal neural dynamics during grasping. Proceedings of the National Academy of Sciences 117, 32124–32135. doi: 10.1073/pnas.2005087117

Montalà-Flaquer, M., López-León, C. F., Tornero, D., Houben, A. M., Fardet, T., Monceau, P., et al. (2022). Rich dynamics and functional organization on topographically designed neuronal networks in vitro. iScience 25, 105680. doi: 10.1016/j.isci.2022.105680

Morrison, A., Diesmann, M., and Gerstner, W. (2008). Phenomenological models of synaptic plasticity based on spike timing. Biol Cybern 98, 459–478. doi: 10.1007/s00422-008-0233-1

Murata, Y., Higo, N., Hayashi, T., Nishimura, Y., Sugiyama, Y., Oishi, T., et al. (2015). Temporal Plasticity Involved in Recovery from Manual Dexterity Deficit after Motor Cortex Lesion in Macaque Monkeys. J. Neurosci. 35, 84–95. doi: 10.1523/JNEUROSCI.1737-14.2015

Nagappan, P. G., Chen, H., and Wang, D.-Y. (2020). Neuroregeneration and plasticity: a review of the physiological mechanisms for achieving functional recovery postinjury. Military Medical Research 7, 30. doi: 10.1186/s40779-020-00259-3

Nicola, W., and Clopath, C. (2017). Supervised learning in spiking neural networks with FORCE training. Nat Commun 8, 2208. doi: 10.1038/s41467-017-01827-3

Nudo, R. J. (2013). Recovery after brain injury: mechanisms and principles. Front Hum Neurosci 7, 887. doi: 10.3389/fnhum.2013.00887

O’Neill, K. M., Anderson, E. D., Mukherjee, S., Gandu, S., McEwan, S. A., Omelchenko, A., et al. (2023). Time-dependent homeostatic mechanisms underlie brain-derived neurotrophic factor action on neural circuitry. Commun Biol 6, 1–26. doi: 10.1038/s42003-023-05638-9

Onnela, J.-P., Saramäki, J., Kertész, J., and Kaski, K. (2005). Intensity and coherence of motifs in weighted complex networks. Phys. Rev. E 71, 065103. doi: 10.1103/PhysRevE.71.065103

Orlandi, J. G., Soriano, J., Alvarez-Lacalle, E., Teller, S., and Casademunt, J. (2013). Noise focusing and the emergence of coherent activity in neuronal cultures. Nature Phys 9, 582–590. doi: 10.1038/nphys2686

Pasquale, V., Martinoia, S., and Chiappalone, M. (2017). Stimulation triggers endogenous activity patterns in cultured cortical networks. Sci Rep 7, 9080. doi: 10.1038/s41598-017-08369-0

Psaier, H., and Dustdar, S. (2011). A survey on self-healing systems: approaches and systems. Computing 91, 43–73. doi: 10.1007/s00607-010-0107-y

Rossum, M. C. W. van, Bi, G. Q., and Turrigiano, G. G. (2000). Stable Hebbian Learning from Spike Timing-Dependent Plasticity. J. Neurosci. 20, 8812–8821. doi: 10.1523/JNEUROSCI.20-23-08812.2000

Rubin, J., Lee, D. D., and Sompolinsky, H. (2001). Equilibrium Properties of Temporally Asymmetric Hebbian Plasticity. Phys. Rev. Lett. 86, 364–367. doi: 10.1103/PhysRevLett.86.364

Rubinov, M., and Sporns, O. (2010). Complex network measures of brain connectivity: uses and interpretations. Neuroimage 52, 1059–1069. doi: 10.1016/j.neuroimage.2009.10.003

Schmidgall, S., Ziaei, R., Achterberg, J., Kirsch, L., Hajiseyedrazi, S. P., and Eshraghian, J. (2024). Brain-inspired learning in artificial neural networks: A review. APL Machine Learning 2, 021501. doi: 10.1063/5.0186054

Slaney, M., and Lyon, R. (1993). On the importance of time—a temporal representation of sound. Available at: https://www.semanticscholar.org/paper/On-the-importance-of-time%E2%80%94a-temporal-representation-Slaney-Lyon/cbfc0cc32d1670a2c5421b2872fa6afadcb796b9 (Accessed November 26, 2024).

Soriano, J. (2023). Neuronal Cultures: Exploring Biophysics, Complex Systems, and Medicine in a Dish. Biophysica 3, 181–202. doi: 10.3390/biophysica3010012

Sporns, O., and Betzel, R. F. (2016). Modular Brain Networks. Annu. Rev. Psychol. 67, 613–640. doi: 10.1146/annurev-psych-122414-033634

Stetter, O., Battaglia, D., Soriano, J., and Geisel, T. (2012). Model-Free Reconstruction of Excitatory Neuronal Connectivity from Calcium Imaging Signals. PLOS Computational Biology 8, e1002653. doi: 10.1371/journal.pcbi.1002653

Sumi, T., Yamamoto, H., and Hirano-Iwata, A. (2020). Suppression of hypersynchronous network activity in cultured cortical neurons using an ultrasoft silicone scaffold. Soft Matter 16, 3195–3202. doi: 10.1039/C9SM02432H

Sumi, T., Yamamoto, H., Katori, Y., Ito, K., Moriya, S., Konno, T., et al. (2023). Biological neurons act as generalization filters in reservoir computing. Proceedings of the National Academy of Sciences 120, e2217008120. doi: 10.1073/pnas.2217008120

Takemuro, T., Yamamoto, H., Sato, S., and Hirano-Iwata, A. (2020). Polydimethylsiloxane microfluidic films for in vitro engineering of small-scale neuronal networks. Jpn. J. Appl. Phys. 59, 117001. doi: 10.35848/1347-4065/abc1ac

Teller, S., Estévez-Priego, E., Granell, C., Tornero, D., Andilla, J., Olarte, O. E., et al. (2020). Spontaneous Functional Recovery after Focal Damage in Neuronal Cultures. eNeuro 7, ENEURO.0254-19.2019. doi: 10.1523/ENEURO.0254-19.2019

Tibau, E., Ludl, A.-A., Rüdiger, S., Orlandi, J. G., and Soriano, J. (2020). Neuronal Spatial Arrangement Shapes Effective Connectivity Traits of in vitro Cortical Networks. IEEE Transactions on Network Science and Engineering 7, 435–448. doi: 10.1109/TNSE.2018.2862919

Turrigiano, G. G. (1999). Homeostatic plasticity in neuronal networks: the more things change, the more they stay the same. Trends in Neurosciences 22, 221–227. doi: 10.1016/S0166-2236(98)01341-1

Turrigiano, G. G. (2008). The Self-Tuning Neuron: Synaptic Scaling of Excitatory Synapses. Cell 135, 422–435. doi: 10.1016/j.cell.2008.10.008

Vishwanathan, A., Sood, A., Wu, J., Ramirez, A. D., Yang, R., Kemnitz, N., et al. (2024). Predicting modular functions and neural coding of behavior from a synaptic wiring diagram. Nat Neurosci 27, 2443–2454. doi: 10.1038/s41593-024-01784-3

Wagenaar, D. A., Pine, J., and Potter, S. M. (2006). An extremely rich repertoire of bursting patterns during the development of cortical cultures. BMC Neurosci 7, 11. doi: 10.1186/1471-2202-7-11

Walker, K. R., and Tesco, G. (2013). Molecular mechanisms of cognitive dysfunction following traumatic brain injury. Front Aging Neurosci 5, 29. doi: 10.3389/fnagi.2013.00029

Wang, C., Lin, C., Zhao, Y., Samantzis, M., Sedlak, P., Sah, P., et al. (2023). 40-Hz optogenetic stimulation rescues functional synaptic plasticity after stroke. Cell Reports 42, 113475. doi: 10.1016/j.celrep.2023.113475

Yada, Y., Yasuda, S., and Takahashi, H. (2021). Physical reservoir computing with FORCE learning in a living neuronal culture. Applied Physics Letters 119, 173701. doi: 10.1063/5.0064771

Yamamoto, H., Kubota, S., Chida, Y., Morita, M., Moriya, S., Akima, H., et al. (2016). Size-dependent regulation of synchronized activity in living neuronal networks. Phys Rev E 94, 012407. doi: 10.1103/PhysRevE.94.012407

Yamamoto, H., Moriya, S., Ide, K., Hayakawa, T., Akima, H., Sato, S., et al. (2018). Impact of modular organization on dynamical richness in cortical networks. Science Advances 4, eaau4914. doi: 10.1126/sciadv.aau4914

Yamamoto, H., Spitzner, F. P., Takemuro, T., Buendía, V., Murota, H., Morante, C., et al. (2023). Modular architecture facilitates noise-driven control of synchrony in neuronal networks. Science Advances 9, eade1755. doi: 10.1126/sciadv.ade1755

Zhang, T., Huang, Q., Jiao, C., Liu, H., Nie, B., Liang, S., et al. (2019). Modular architecture of metabolic brain network and its effects on the spread of perturbation impact. NeuroImage 186, 146–154. doi: 10.1016/j.neuroimage.2018.11.003

