## Supplementary Figures for "Modular architecture confers robustness to damage and facilitates recovery in spiking neural networks modeling *in vitro* neurons"

**This PDF file includes:**

Figures S1 to S8

### Supplementary Figures

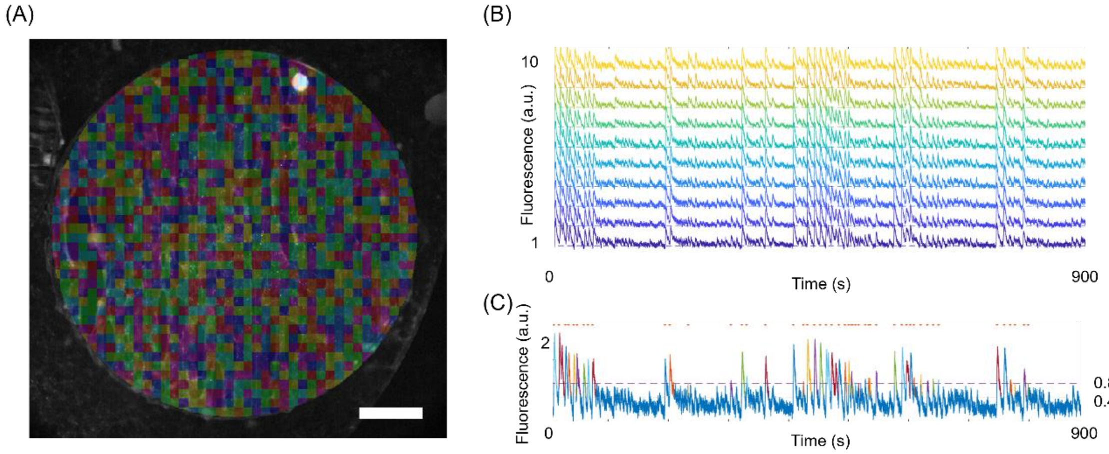


**Supplementary Figure S1. (A)** Selection of Region of Interest (ROI). A 150 µm × 150 µm square region of calcium intensity data was designated as the ROI for neurons cultured on 6 mm diameter PDMS disks. Approximately 1,280 ROIs were selected. Scale bar is 1 mm. **(B)** Representative mean fluorescence intensity data in the ROIs. Ten representative examples were plotted. **(C)** Spike detection using a Schmitt trigger (lower threshold = 0.4, upper threshold = 0.8). Spikes are indicated by orange dots in the plot.

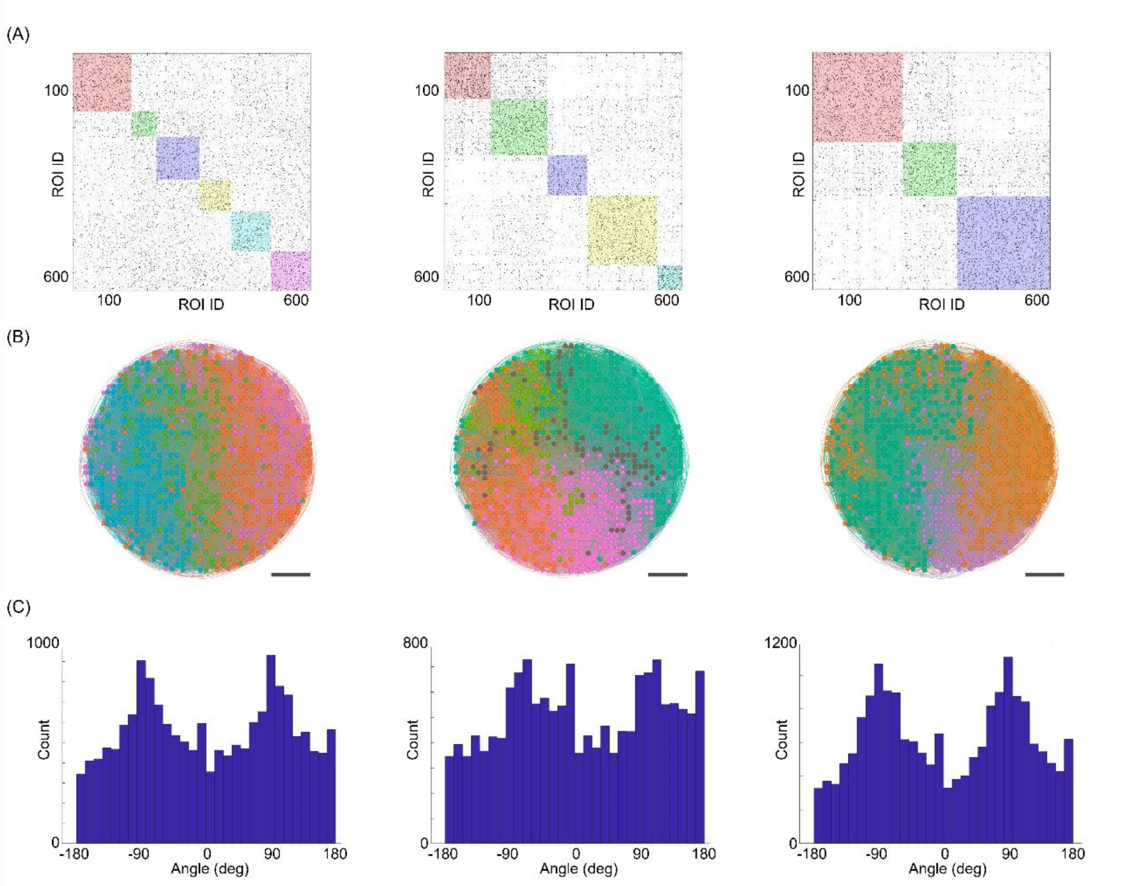


**Supplementary Figure S2. (A)** Effective connectivity maps estimated by transfer entropy before (left), immediately after (center), and 24 hours after (right) the injury. The colors represent modules when modularity is maximized. To enhance visualization, the analysis includes half the total ROIs. **(B)** Visualization of effective connectivity and modules. Vertical connections along the underlying PDMS topographical tracks form distinct modules (left), which are temporarily disorganized due to the injury (center). However, after 24 hours, both the vertical connections and modules re-form (right). Scale bars are 1 mm. **(C)** Histogram of connection angles. Before the injury, sharp peaks appear at -90° and 90° (left), reflecting the dominance of vertical connections. These peaks are blunted after injury (center), but reappear 24 hours later (right).

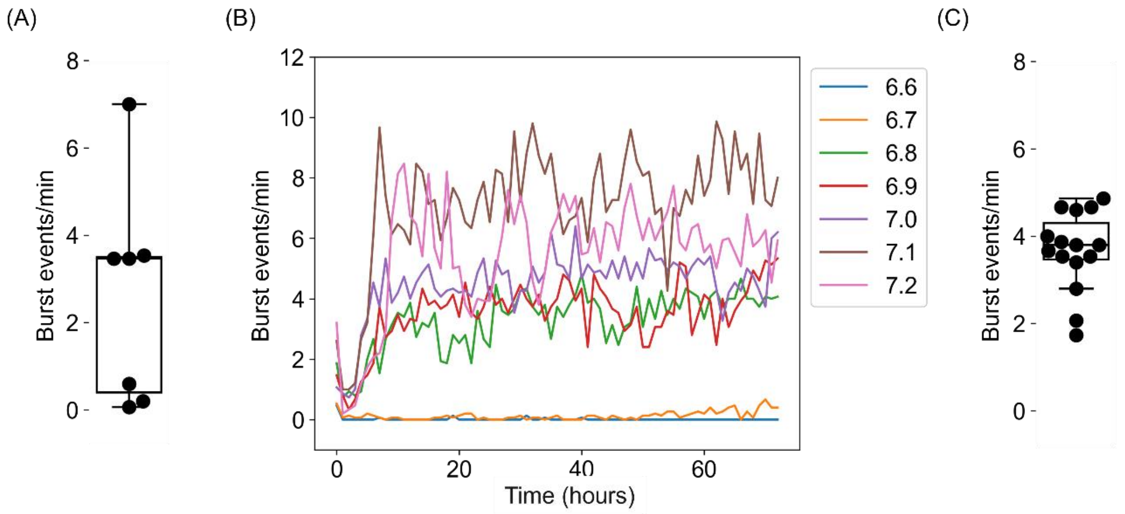


**Supplementary Figure S3. (A)** Burst event rate per minute in cultured neuronal networks before the injury. Due to the wide variability, the median value of 3.47 was chosen as the target for parameter adjustment in the simulation. **(B)** Burst event rate for different maximum synaptic weights ($w_{max}$). The event rate initially increases due to STDP but stabilizes after approximately 30 simulated hours. As $w_{max}$ increases, the final event rate rises. The value $w_{max}$ = 6.8 was selected as it most closely matched the target rate of 3.47. **(C)** Event rate after 72 simulated hours of STDP at $w_{max}$ = 6.8. The resulting rate was 3.67 ± 0.88 (mean ± SD), closely matching the value observed in the cultured neuronal network.

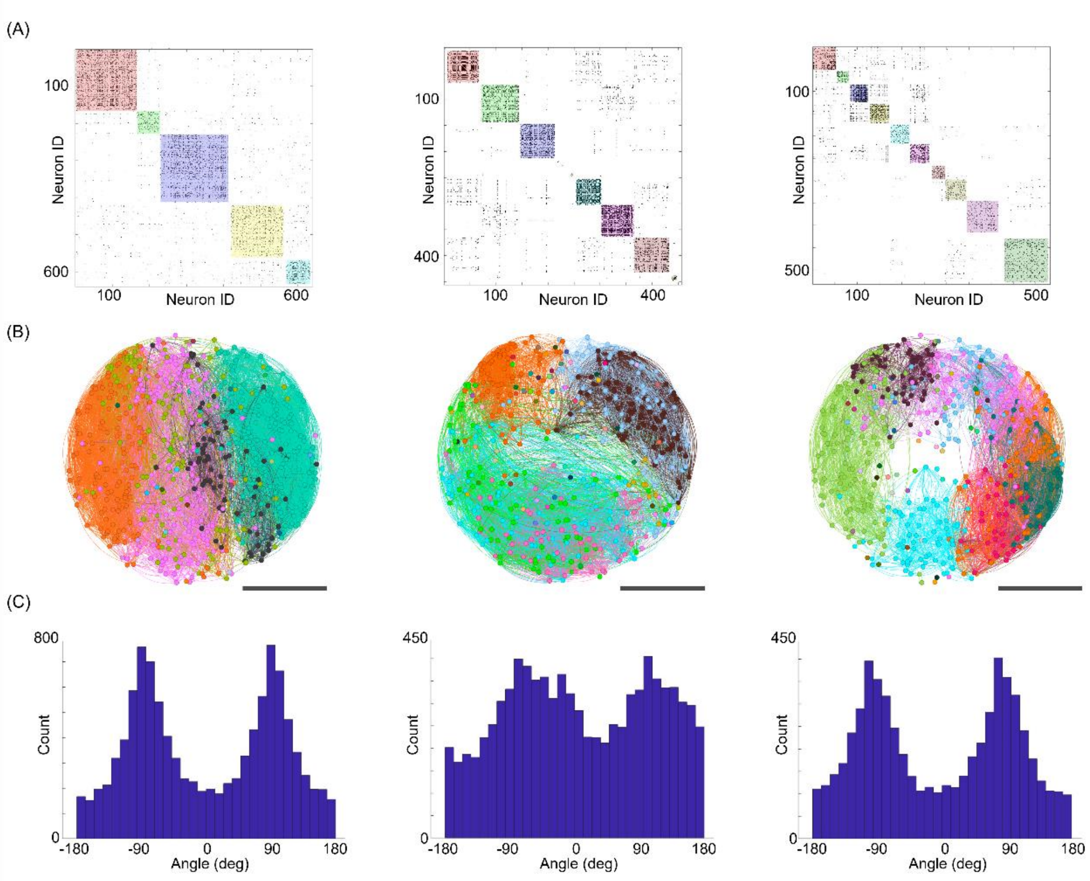


**Supplementary Figure S4. (A)** Effective connectivity in the spiking neural network model before (left), immediately after (center), and 24 hours after (right) the injury. The colors represent modules when modularity is maximized. To improve visualization, the analysis includes one-quarter of the total neurons. **(B)** Visualization of effective connectivity and modules. Similar to the culture experiment, vertical connections along the track pattern form distinct modules (left), which are temporarily disorganized following the injury (center). After 24 hours, the vertical connections and modules are formed again (right). **(C)** Histogram of connection angles. As in the culture experiment, sharp peaks at -90° and 90° are observed before the injury (left). These peaks blunted after the injury (center) but reappear after 24 hours (right)

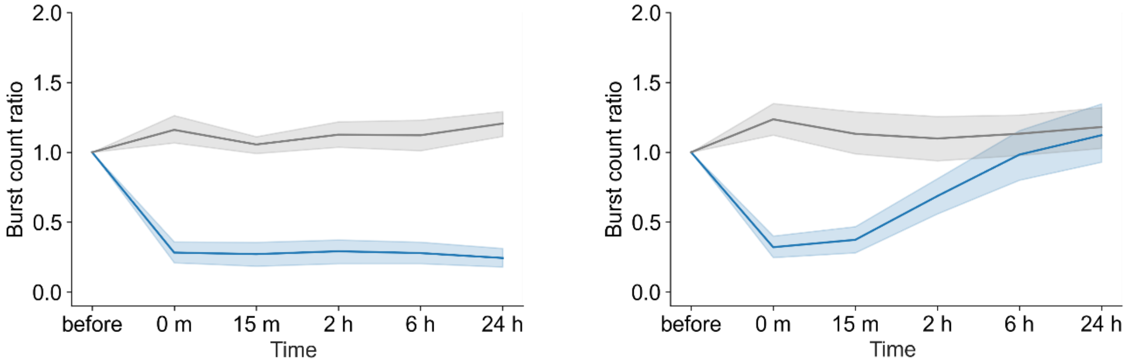


**Supplementary Figure S5.** Time course of burst event rate without STDP (left) and with STDP (right). The blue line shows damaged samples, and the grey line shows undamaged controls. Shaded areas represent the 95% confidence interval. In undamaged samples, burst activity remained stable over time. Damage initially reduced activity, followed by sustained low activity without STDP. However, with STDP, the activity recovered to a level comparable to the undamaged samples within 24 hours.

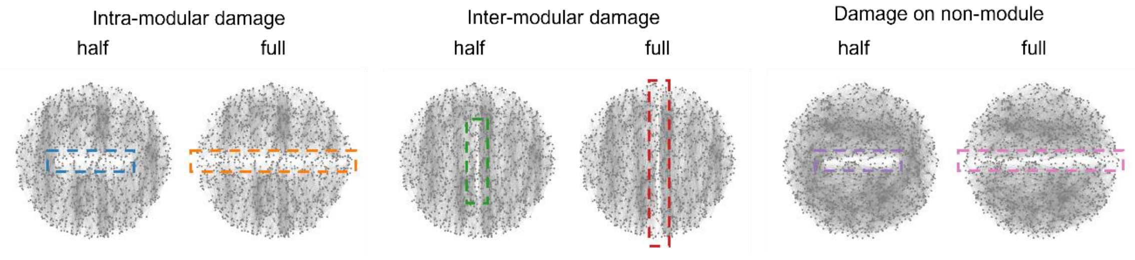


**Supplementary Figure S6.** Representative network of damage actions on patterned (modular) and homogeneous networks. Intra-modular, inter-modular damages, and Damage on non-module as well as the definitions of 'half' and 'full' are described in Figure 4A

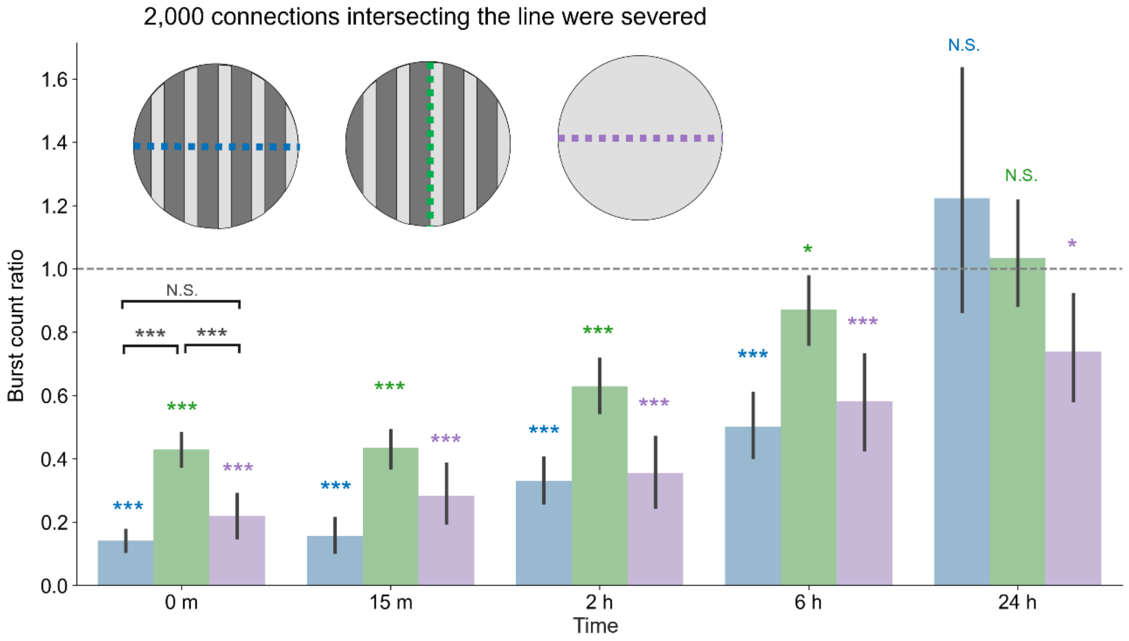


**Supplementary Figure S7.** Burst frequency changes for scenarios with the same number of connections to be disconnected. For each scenario, the color bar shows the ratio of burst frequency between the post- and pre-injury conditions. The bar height indicates the mean, and error bars indicate 95% confidence intervals. Color asterisks indicate a two-sided 1-sample *t*-test (*$p < 0.05$; **$p < 0.01$; ***$p < 0.001$; N.S., no significance; $n = 15$, $df = 14$). Asterisks in black correspond to a two-sided unpaired *t*-test (**$p < 0.01$; ***$p < 0.001$; N.S., no significance; $n = 15$, $df = 13$). Even when the number of disconnections was common, the effects of the direction of damage and the background module structure were consistent with the results in Figure 4B.

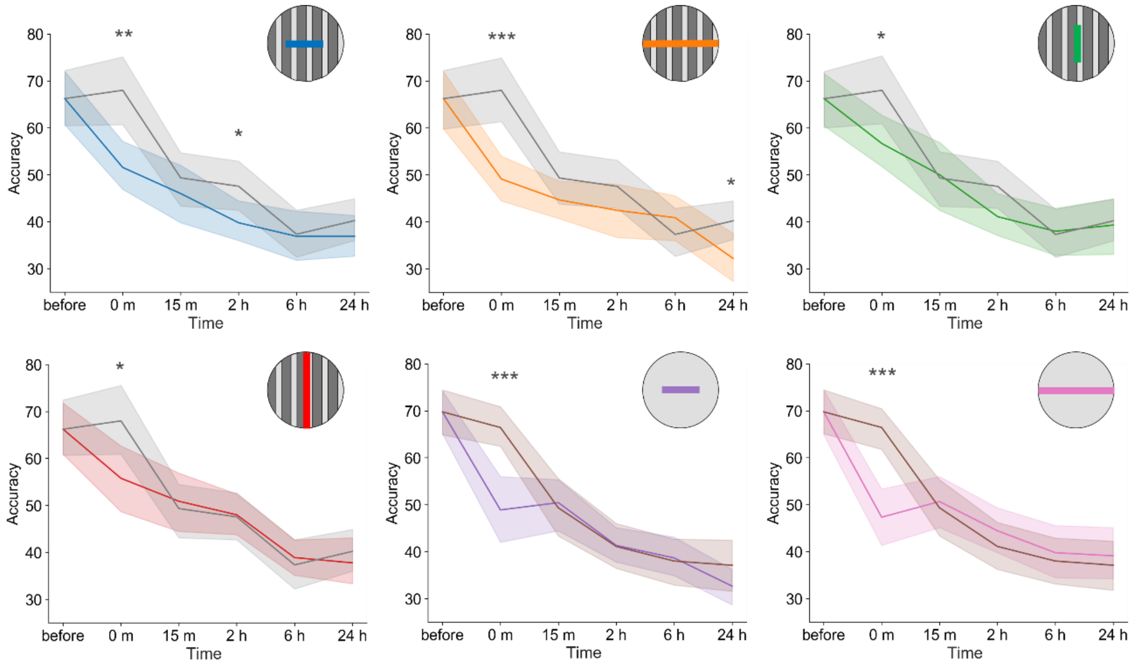


**Supplementary Figure S8.** Time courses of accuracy in the reservoir computing tasks for the six damaging conditions. The colors correspond to each damaging action. The training of the output layer stopped before the damage. Grey curves show the undamaged case for the tracks-patterned network, and the brown curves show the undamaged case for the control, unpatterned network. In the panels, the lines indicate the averaged value, and shadings the 95% confidence interval. *$p$ < 0.05; **$p$ < 0.01; ***$p$ < 0.001 (two-sided unpaired $t$-test, $n$ = 15, $df$ = 13).
